# Advancing Protein Ensemble Predictions Across the Order–Disorder Continuum

**DOI:** 10.1101/2025.10.18.680935

**Authors:** Michele Invernizzi, Sandro Bottaro, Julian O. Streit, Bruno Trentini, Niccolò Alberto Elia Venanzi, Danny Reidenbach, Youhan Lee, Christian Dallago, Hassan Sirelkhatim, Bowen Jing, Fabio Airoldi, Kresten Lindorff-Larsen, Carlo Fisicaro, Kamil Tamiola

## Abstract

While deep learning has transformed structure prediction for ordered proteins, intrinsically disordered proteins remain poorly predicted due to systematic underrepresentation in training data, despite constituting approximately 30% of eukaryotic proteomes. We introduce PeptoneBench, the first benchmark to enable systematic assessment of ensemble generators for both ordered and disordered proteins, integrating diverse experimental observables. Our analysis reveals that existing evaluation metrics exhibit systematic bias toward the structured spectrum of the proteome. Assessment of popular predictors (AlphaFold2, ESMFlow, Boltz2) confirms high accuracy on ordered proteins but shows performance degradation with increasing disorder. We further present PepTron, a flow-matching ensemble generator trained on data augmented with synthetic disordered protein ensembles. On our benchmark PepTron matches BioEmu on disordered regions while maintaining competitive accuracy on ordered protein benchmarks. Our data augmentation approach demonstrates that targeted training strategies can approach the performance of computationally expensive simulation-based methods, establishing a generalizable framework applicable to other protein generative models. All datasets, models, and code are openly available.

## 1 Introduction

Intrinsically disordered proteins (IDPs) and intrinsically disordered regions (IDRs), which lack stable tertiary structure under native conditions, play central roles in signaling, transcriptional regulation, and a wide range of human diseases [1]. Computational analyses estimate that approximately 30% of residues in eukaryotic proteomes are disordered, yet these regions remain difficult to characterize experimentally and computationally [2].

The flexible nature of IDPs complicates analysis by traditional structure determination methods such as X-ray crystallography and cryogenic sample electron microscopy, necessitating ensemble-averaged approaches like nuclear magnetic resonance (NMR) spectroscopy and small-angle X-ray scattering (SAXS), and other biophysical techniques. These experimental limitations, in turn, influence computational modeling: modern protein structure prediction methods inherit a bias toward ordered structures from their reliance on the Protein Data Bank (PDB) [3, 4], where disordered regions are systematically underrepresented [5, 6]. Proteome-wide analyses consistently report substantially higher fractions of disordered residues than appear in structural databases [2, 7]. Furthermore, this bias is compounded by poor sequence conservation and sparse multiple sequence alignments (MSAs) of IDRs, limiting the effectiveness of MSA-driven features for disorder modeling [8]. Consequently, models such as AlphaFold2, which are trained on the PDB, struggle to generalize to the substantial fraction of proteins that contain IDRs [9].

Recent methodological advances have begun to address these limitations. While AlphaFold2’s predictions for disordered regions do not fully capture biophysical reality, its per-residue local distance difference test (pLDDT) score is remarkably effective for disorder identification [10]. AlphaFold3 employs cross-distillation using AlphaFold-Multimer predictions as training signals to represent disordered regions as extended structures, thereby mitigating hallucinations [11]. Similarly, RosettaFold-3 adopts artificially extended backbone representations for disordered regions during training [12]. Beyond single-structure predictors, ensemble generation methods directly emulate protein dynamics. AlphaFlow and ESM-Flow [13] combine flow-matching frameworks with structure prediction models to generate conformational ensembles, while BioEmu [14] integrates over 200 milliseconds of molecular dynamics simulations alongside ordered structures and experimental stability data at training stage to generate protein ensembles at scale.

Complementing deep learning approaches, physics-based methods offer alternative strategies to address IDPs[15, 16], while others utilize generative adversarial or diffusion models trained on molecular simulations [17, 18]. Unified predictors capable of capturing both ordered and disordered regimes have also emerged [19–21].

Regardless of the predictive method employed, evaluating predictions for disordered proteins presents inherent difficulties: no single native structure exists, and metrics such as root-mean-square deviation (RMSD) and global distance test (GDT-TS) are ill-posed. Meaningful validation requires ensemble-oriented approaches that compare predicted conformational ensembles to experimentally accessible observables: chemical shifts (CS), SAXS profiles, residual dipolar couplings (RDCs), and paramagnetic relaxation enhancements (PREs) [22–24], aligning deep-learning modeling with practices long established in force-field development [25–27].

In this work, we introduce two principal contributions to address these gaps:

- **Evaluation framework:** We introduce PeptoneBench, a systematic protocol with accompanying code for ensemble evaluation that compares experimental observables to ensemble-averaged predictions, including maximum entropy reweighting [28–31], leveraging a curated dataset of SAXS, NMR, RDC, and PRE data.
- **Model development:** We present PepTron, a generative model trained on an augmented dataset enriched with synthetic IDR ensembles. PepTron is benchmarked against leading predictors (AlphaFold2, Boltz2, BioEmu) via our framework, demonstrating state-of-the-art consistency with experimental ensemble properties across both ordered and disordered regimes.

Together, these contributions establish a promising framework for more accurate, ensemble-based protein structure prediction, improving representation of both ordered and disordered configurations. All datasets, models, and code are available under the CC BY or Apache 2.0 license, as detailed in Sec. 3.

## 2 Results

### 2.1 PeptoneBench: Disorder-aware Protein Ensembles Evaluation

The Protein Data Bank (PDB) [3] is foundational for training and validating deep learning models in protein structure prediction. However, it is notably depleted in IDRs and IDPs, introducing a bias toward ordered structures (Fig. 1a). Our analysis indicates that only ∼ 3% of PDB entries are classified as disordered, compared to 20–30% in the human proteome, consistent with previous estimates [32, 33]. This underrepresentation motivates the use of alternative databases: the IDRome database [34], designed to encompass all disordered regions in the human proteome, is highly enriched in disorder (approximately 77%), providing complementary coverage. However, experimentally determined protein structures for the IDRome are not available.

**Fig. 1.**
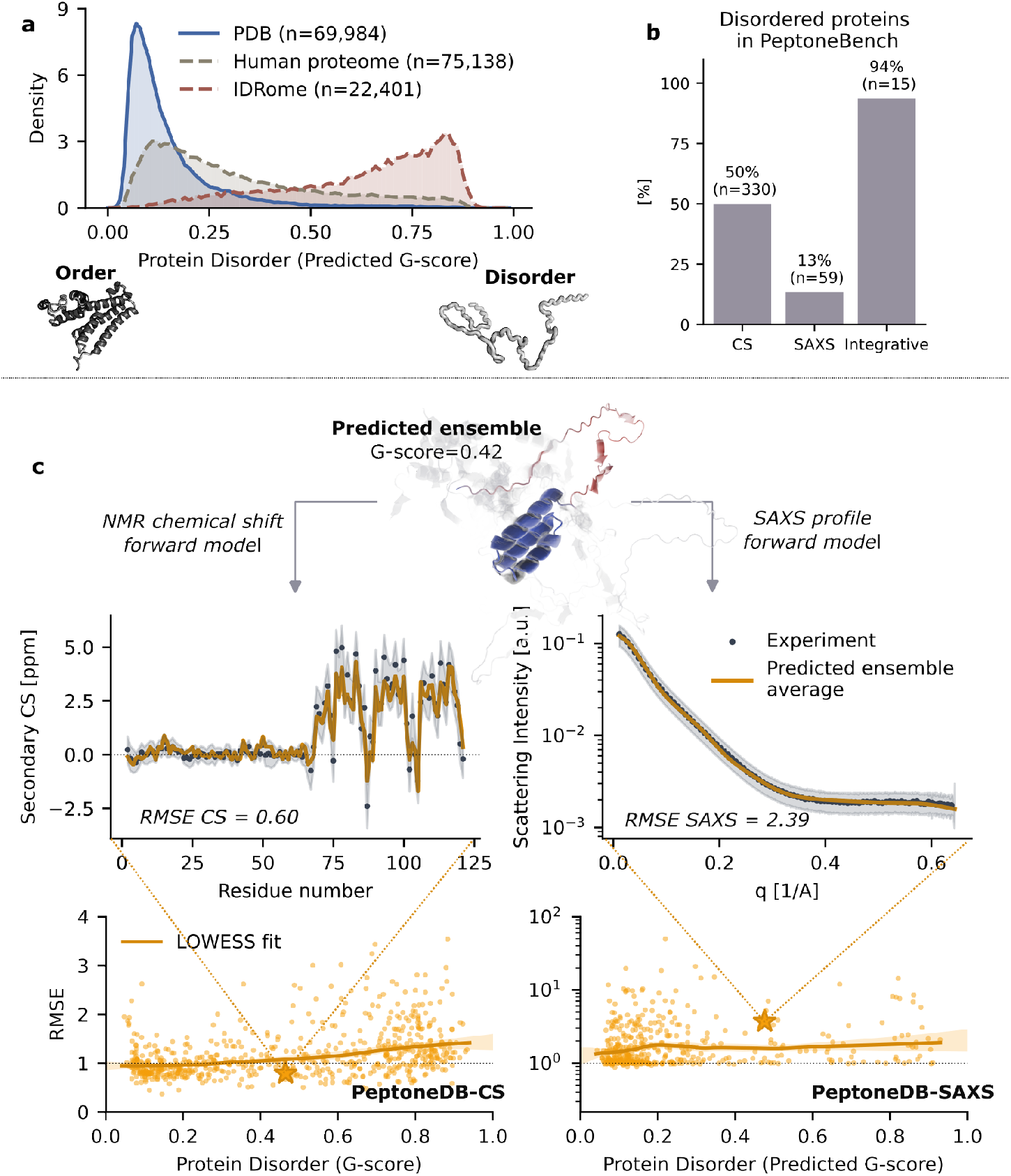
(**a**) Distribution of predicted protein disorder (mean G-score) in the PDB, human proteome, and IDRome datasets, with examples of ordered and disordered structures. Each reported dataset was clustered with MMseqs2, with sizes indicated in parentheses. (**b**) Fraction of disordered proteins (average G-score *>* 0.5) in PeptoneBench databases. (**c**) Overview of the benchmarking workflow. For each protein sequence, an ensemble is predicted using the chosen model (in this case PepTron), enabling calculation of relevant observables (e.g., secondary CS, SAXS profiles) depending on the dataset. Ensemble-averaged predictions (with or without maximum entropy reweighting) are compared to experimental measurements, and uncertainty-normalized RMSE is computed for each PeptoneDB-CS and PeptoneDB-SAXS entry. Normalized RMSE values are visualized as a function of disorder content (G-score), with data aggregated via LOWESS smoothing and uncertainty estimated by bootstrapping. Stars denote the same protein present in both PeptoneDB-CS and PeptoneDB-SAXS datasets.

Disorder is quantified here using the protein mean G-score, which ranges from 0 (fully ordered) to 1 (fully disordered) [35]. G-scores are derived from secondary chemical shifts (CS), which directly report on the presence or absence of local secondary structure [36]. For proteins lacking experimental CS data, we predict G-scores using ADOPT2 [37], a machine learning framework trained on the TriZOD dataset [35].

Quantifying disorder makes it clear that, in order to evaluate the quality of generated protein ensembles, one has to go beyond the PDB, and look for different sources of experimental validation. To this end we introduce three curated datasets PeptoneDB-CS (NMR chemical shifts from BMRB [38]), PeptoneDB-SAXS (SAXS profiles from SASBDB [39]), and PeptoneDB-Integrative (entries with multiple complementary experimental measurements [15, 26, 40]; see Table 1) and Fig. 1b. These datasets cover the whole order-disorder continuum and provide complementary information: CS strongly correlate with local secondary structure content, while SAXS curves are related to the overall shape of the protein ensemble. The PeptoneDB-Integrative focuses on IDPs and provides for each entry an orthogonal measurement (RDCs for CS and PREs for SAXS, see Table A1) that can be used for cross-validation.

**Table 1.** Curated protein datasets used in this study, showing data sources and counts of ordered and disordered proteins and residues. Proteins are classified as ordered (mean G-score *<* 0.5) or disordered (mean G-score *>* 0.5). For PeptoneDB-SAXS, G-scores are predicted when chemical shift data are unavailable.*Note:* PaaA2 in PeptoneDB-Integrative (mean G-score = 0.43; see Table A1) is classified as ordered by this threshold, but is commonly reported as disordered in the literature [45].

| Dataset | Data type | Source | # proteins<br>(ordered/disordered) | # residues<br>(ordered/disordered) |
| --- | --- | --- | --- | --- |
| PeptoneDB-CS | NMR chemical shifts | BMRB | 329 / 330 | 32,908 / 31,507 |
| PeptoneDB-SAXS | SAXS profiles | SASBDB | 380 / 59 | 92,577 / 18,372 |
| PeptoneDB-Integrative | NMR chemical shifts<br>RDCs, SAXS, PREs | Literature | 1 / 15 | 151 / 2,042 |

We use these datasets to implement *PeptoneBench*, a systematic evaluation framework for quantifying predicted protein ensembles agreement with experimental data (Fig. 1c), across the order-disorder continuum. Prediction models generate ensembles that are pre-processed using standardized criteria (Appendix A.1). Experimental observables are computed from predicted structures using appropriate forward models (Appendix A.2, A.3, A.4) [41]. Agreement between prediction and experiment is assessed using normalized root mean squared error (RMSE), Eq. A6, accounting for uncertainties from both forward models and experimental data [42–44]. RMSE values are visualized as a function of protein disorder (G-score), with data aggregated via locally weighted scatterplot smoothing (LOWESS) and uncertainty estimated by bootstrapping. The area under the LOWESS curves (AUC) provides a condensed accuracy metric across the disorder content. For convenience, we define a PeptoneBench summary score by combining the AUC for CS and SAXS, see Eq. D12.

While RMSE is indicative of accuracy, it is insufficient on its own: broader ensembles with higher RMSE may better capture experimental distributions if improved by maximum entropy reweighting [28, 31]. Therefore, PeptoneBench also computes RMSE with post-refinement results, to determine not only how well ensembles match experiments initially, but also how much their agreement can be improved through maximum entropy reweighting. This approach distinguishes between ensembles that contain the right conformations but need better weighting (*reweightable errors*), and those that lack important conformations entirely (*irrecoverable deficiencies*). A description of the robust reweighting procedure developed for PeptoneBench is provided in the Appendix A.5.

Ensemble refinement is particularly relevant when dealing with IDP ensembles, which are highly sensitive to experimental conditions (e.g., temperature, pH, ionic strength). Generative models that sample diverse conformations across a range of conditions, similarly to multicanonical approaches in molecular dynamics, could be reweighted to match specific experimental parameters, greatly enhancing their predictive utility.

### 2.2 PepTron: Augmenting Training Data with Disordered Regions for Improved Protein Ensemble Prediction

We introduce *PepTron*, a protein structure generator based on the flow-matching ESMFlow architecture [13], and designed to generate diverse conformational ensembles for proteins spanning from ordered to disordered states.

PepTron is fine-tuned on both, experimentally determined structures from the PDB, and synthetically generated ensembles of disordered proteins. This augmentation exposes the model to the full order-disorder continuum, enabling more accurate predictions across diverse protein types. Remarkably, even computationally inexpensive methods for generating disordered ensembles substantially improve performance. Performance was assessed using PeptoneBench, ensuring evaluation on experimental data fully withheld from training. We benchmarked PepTron together with ESMFold, ESMFlow, AlphaFold2, Boltz2, and BioEmu, and performed dedicated evaluations on the IDP-focused PeptoneDB-Integrative dataset.

#### 2.2.1 Synthetic IDP Dataset Generation

To address the lack of structural data for disordered regions, we generate a synthetic structural dataset, called IDRome-o, from the sequences in IDRome [46] (Fig. 1a). To this end, we developed a fragment-based protein structure generator, *IDP-o*, which can generate physically plausible IDP ensembles at scale and with reduced computational cost. IDP-o harnesses principles from fragment-based approaches [47] and hierarchical chain-growth techniques [48]. Specifically, it constructs protein configurations by aligning smaller fragments taken from a large structure database. Contrary to previous methods which used the PDB as source for the fragments, IDP-o leverages the full AlphaFold Protein Structure Database (214 million predicted structures), which is orders of magnitude larger. This allows IDP-o to use larger six-residue fragments, and better capture transient helicity content, as shown in the comparison with other IDP-specialised models in Appendix B).

It is important to note that by design IDP-o does not sample a specific equilibrium ensemble distribution, but rather it produces diverse conformational ensembles which represent plausible configurations for a given sequence. Accordingly, these ensembles are most effective when combined with reweighting strategies, or as starting points for further molecular dynamics simulations [48].

#### 2.2.2 Model Architecture and Training Mixture

PepTron is based on ESMFlow architecture, but was implemented within the NVIDIA BioNeMo framework, for improved computational performances [49]. It integrates cuEquivariance triangle attention and uses the Modular Co-Design BioNeMo sub-package for implementing flow matching, as described in Appendix C. Both at training and inference time PepTron uses BioNeMo distributed best practices, leveraging Megatron-style [50] tensor-pipeline-context-expert parallelism, mixed precision, activation checkpointing, and optimized input pipelines to scale efficiently across multi-GPU systems. At inference, PepTron runs from a single checkpoint without MSAs or external ESM weights [51] and directly generates structure ensembles.

We start from the ESMFold weights and retrain the flow matching module using the PDB, as described in the ESMFlow paper [13], but with a twice longer crop length of 512. We then fine-tune PepTron using a mixture of PDB and IDRome-o data, as described in detail in Appendix C.

### 2.3 Comparative Benchmarking

To assess the performance improvement of PepTron over the baseline models ESMFlow and IDP-o, we predicted ensembles for all sequences in the PeptoneDB-CS and PeptoneDB-SAXS datasets, and computed the normalized RMSE following the PeptoneBench procedure. Figure 2b summarizes the agreement as a function of protein disorder content on the PeptoneDB-CS set for ESMFlow, IDP-o, and PepTron. For comparison, we also include single-structure predictions from ESMFold [52].

**Fig. 2.**
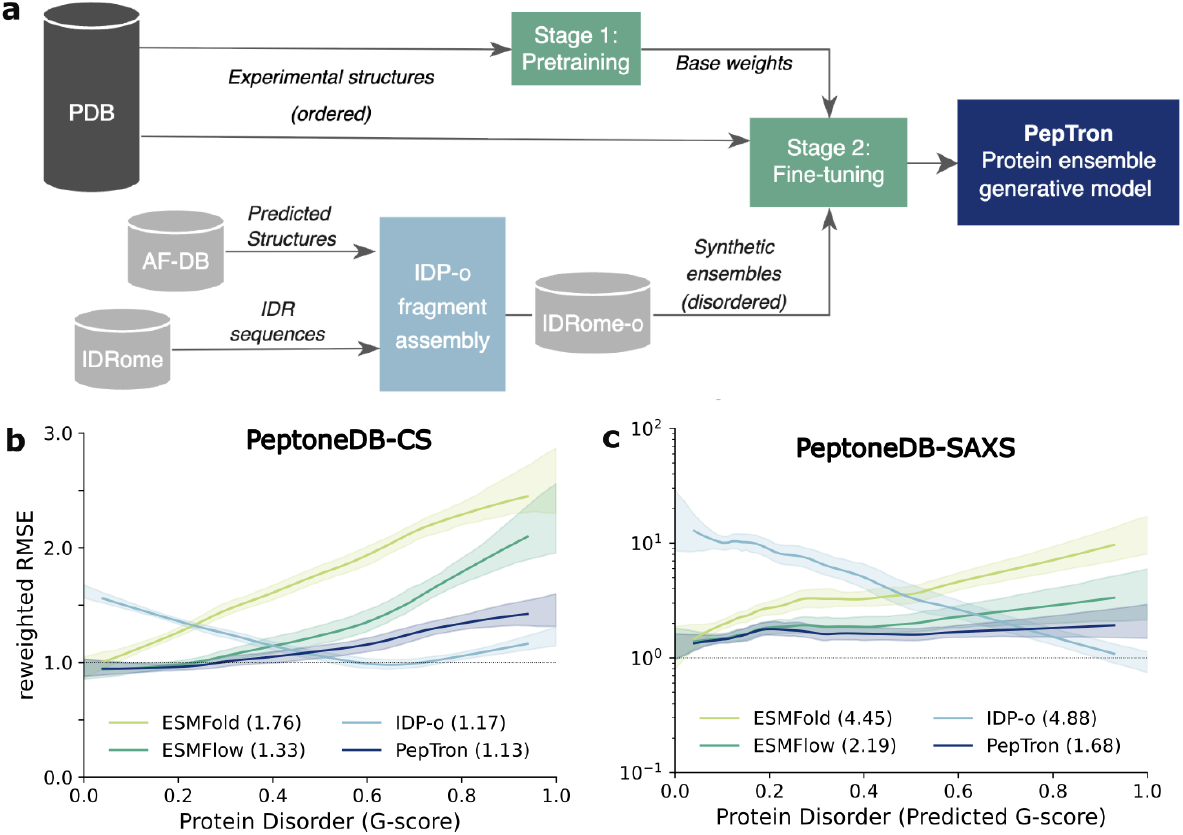
(**a**) Schematic overview of the PepTron model. Structural data from the PDB are used to pretrain the PepTron model which employs the ESMFlow architecture. Synthetic datasets of disordered proteins are generated using IDP-o, drawing sequences from the IDRome and fragments from the AlphaFold UniProt Database (AlphaFold DB). The resulting synthetic dataset is jointly used for fine-tuning, integrating ordered and disordered states. (**b**) Agreement with experimental CS data and (**c**) SAXS profiles. Reported values correspond to per-protein reweighted RMSE fitted via LOWESS regression with bootstrap-estimated uncertainties for ESMFold, ESMFlow, IDP-o, and PepTron. For each model and dataset, we report in parentheses the area under the LOWESS curve as an overall score (lower is better).

A consistent trend emerges: both ESMFold and ESMFlow yield accurate predictions for ordered systems (low G-score, normalized RMSE ≈ 1), but their accuracy declines as disorder increases, with ESM-Fold degrading more rapidly. In contrast, IDP-o displays the opposite pattern, with improved accuracy at higher disorder levels, in line with expectations. PepTron maintains more stable performance across both ordered and disordered protein ensembles. Similar trends are observed when comparing the models against experimental SAXS profiles and upon reweighting (Fig. 2c). This balanced performance suggests that training on diverse synthetic disordered ensembles enables the model to capture conformational heterogeneity without sacrificing accuracy on well-folded structures.

To contextualize the performance of PepTron model on PeptoneBench, we compare it against some of the most widely used protein generative models: AlphaFold2 [11], Boltz2 [53], and BioEmu [14]. Figure 3 shows the overall agreement of these models with experimental CS and SAXS data. For convenience, we also provide the PeptoneBench summary scores in Table 2. Consistent with previous findings for ESM-Fold, both Boltz2 and AlphaFold2 demonstrate strong predictive accuracy for ordered protein systems (low G-score), yet their performance deteriorates as disorder content increases. This systematic decrease highlights a key limitation of current leading models, which, despite their groundbreaking successes in structure prediction, remain challenged by the inherent heterogeneity of IDPs and proteins featuring significant disordered regions.

**Table 2.**
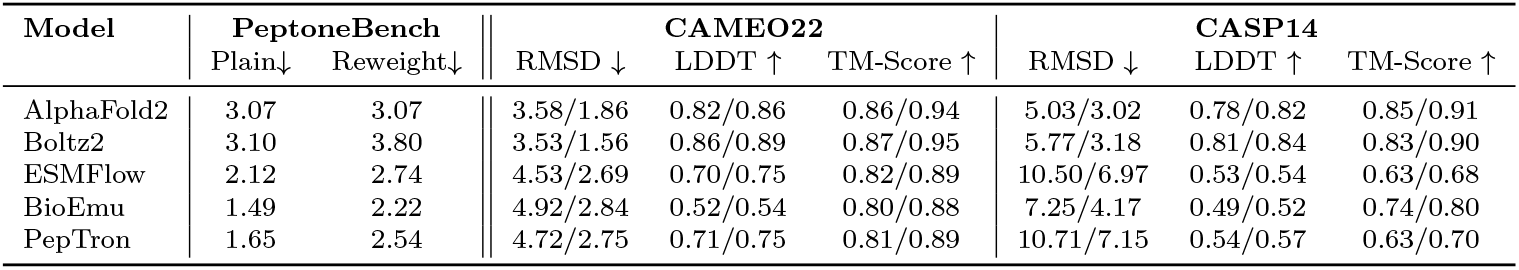
Performance of protein structure prediction on PeptoneBench and the CAMEO22 and CASP14 benchmarks. For the PeptoneBench we report summary scores (see Eq. D12) both with and without ensemble refinement via maximum entropy reweighting. For CAMEO22 and CASP14 we report the average and median over samples for each metric (values for ESMFlow and AlphaFold2 are taken from Ref. [54]).

**Fig. 3.**
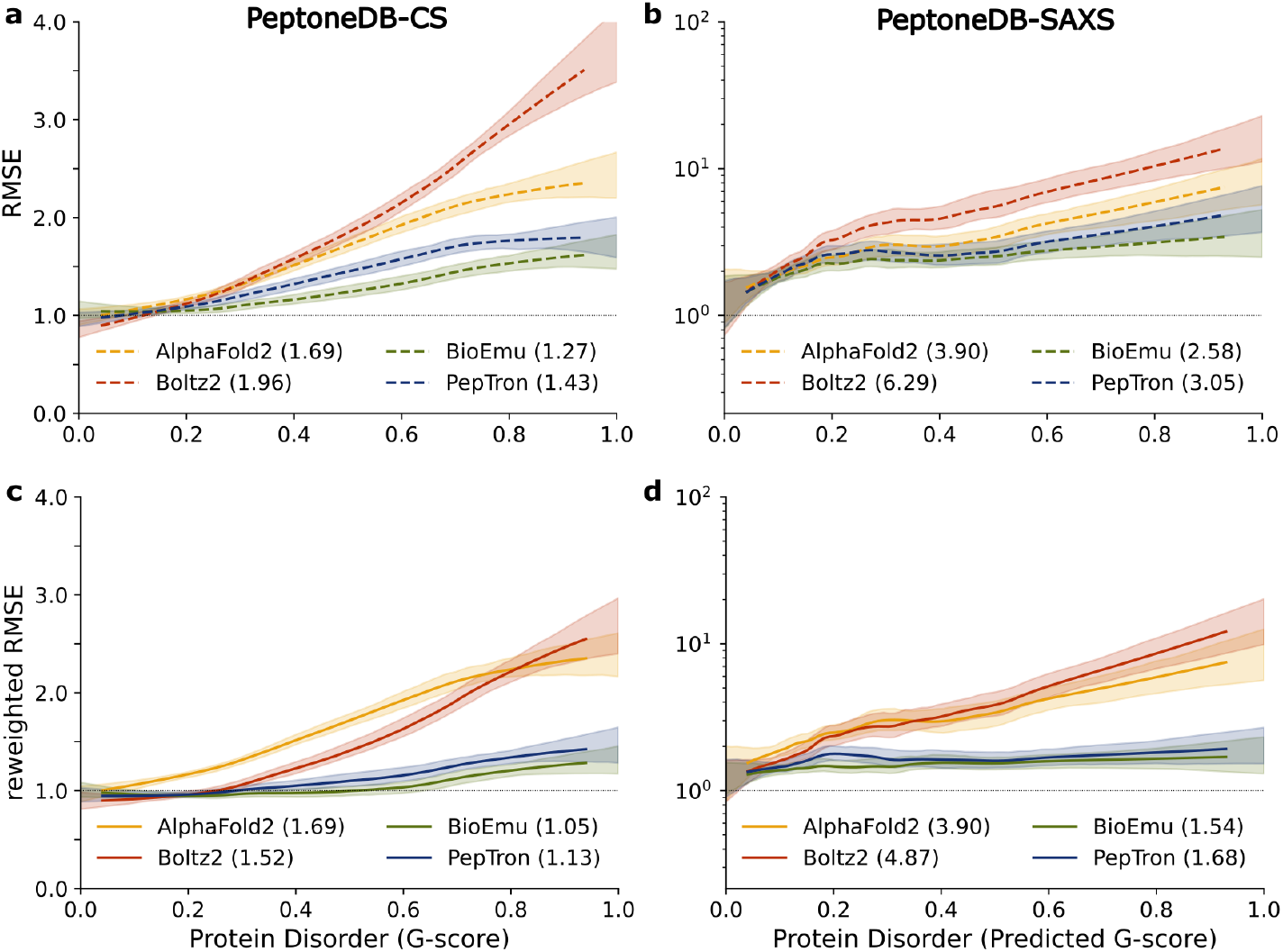
Agreement with experimental CS data (**a**) and SAXS profiles (**b**). Reported values correspond to per-protein normalized RMSE fitted via LOWESS regression with bootstrap-estimated uncertainties for PepTron, AlphaFold2, Boltz2, and BioEmu. For each model and dataset, we report in parentheses the area under the LOWESS curve as a summary score (lower is better). Panels **c** and **d** report the same quantities after maximum entropy reweighting.

By contrast, PepTron and BioEmu exhibit robust performance across the entire order-disorder spectrum, maintaining comparable levels of accuracy for both ordered and disordered proteins. The consistently high agreement observed for BioEmu aligns with its design, as the model has been trained explicitly on extensive molecular dynamics simulations of unfolded proteins that provide rich coverage of conformational diversity well suited to the characteristics of IDPs. This observed trend is consistent across both NMR and SAXS benchmark datasets, highlighting its reliability and generality. A more comprehensive comparative analysis including expanded benchmarking across models, both before and after maximum-entropy reweighting, is provided in the Appendix (Figs. D8, D9, and D10). For convenience, we also provide the PeptoneBench summary scores in Table 2. The PeptoneBench results on a larger set of and Fig. D13.

We further benchmarked the predictors on the canonical ordered datasets CAMEO22 and CASP14 (Table 2). AlphaFold2 and Boltz2 consistently achieve the lowest RMSD values and highest LDDT and TM-scores, establishing them as the top-performing methods for modeling ordered proteins. PepTron and ESMFlow display very similar results, with PepTron matching ESMFlow in both RMSD and accuracy metrics across all benchmarks. This parity indicates that the extensive retraining and fine-tuning applied to PepTron for disordered protein ensembles does not compromise its performance on ordered structures, supporting the effectiveness of our IDR augmentation strategy. While BioEmu lags slightly behind PepTron and ESMFlow on the CAMEO22 benchmark, it shows improved performance on CASP14, with TM-scores and RMSD values closer to the best-performing models, AlphaFold2 and Boltz2.

### 2.4 PeptoneDB-Integrative Insights

While PeptoneDB-CS and PeptoneDB-SAXS provide broad coverage of proteins across the order-disorder continuum, validation on IDPs with multiple orthogonal experimental measurements (PeptoneDB-Integrative) offers a more stringent test of ensemble quality. For this dataset, we performed two complementary analyses. First, we performed maximum entropy fitting of CS data, tuning the regularization parameter to avoid overfitting, ensuring that improving agreement with CS data did not degrade agreement with the orthogonal RDC measurements (Fig. D15). Note that for RDCs, the quality factor (Q-factor) is used to quantify performance (see Appendix A.4).

While the modest number of proteins in this dataset limits definitive ranking, some clear trends emerged (Fig. 4). The disorder-specialized predictor, IDP-o, consistently outperformed all other models after reweighting, yielding lower RMSE and Q-factor values and thus reflecting better agreement with experiment. Closer inspection indicates that models with relatively high RMSE and Q-factors (2–3 and *>*1, respectively) often fail to capture key helical propensities, as confirmed by per-protein, per-model analyses (Figs. D17 and D18). Notably, both PepTron and BioEmu exhibited similar performance on RDCs (mean Q-factor ∼ 0.8), although BioEmu performed better on chemical shifts. In contrast, AlphaFold2, ESMFold, and Boltz2 demonstrated consistently poor agreement across both local experimental metrics.

**Fig. 4.**
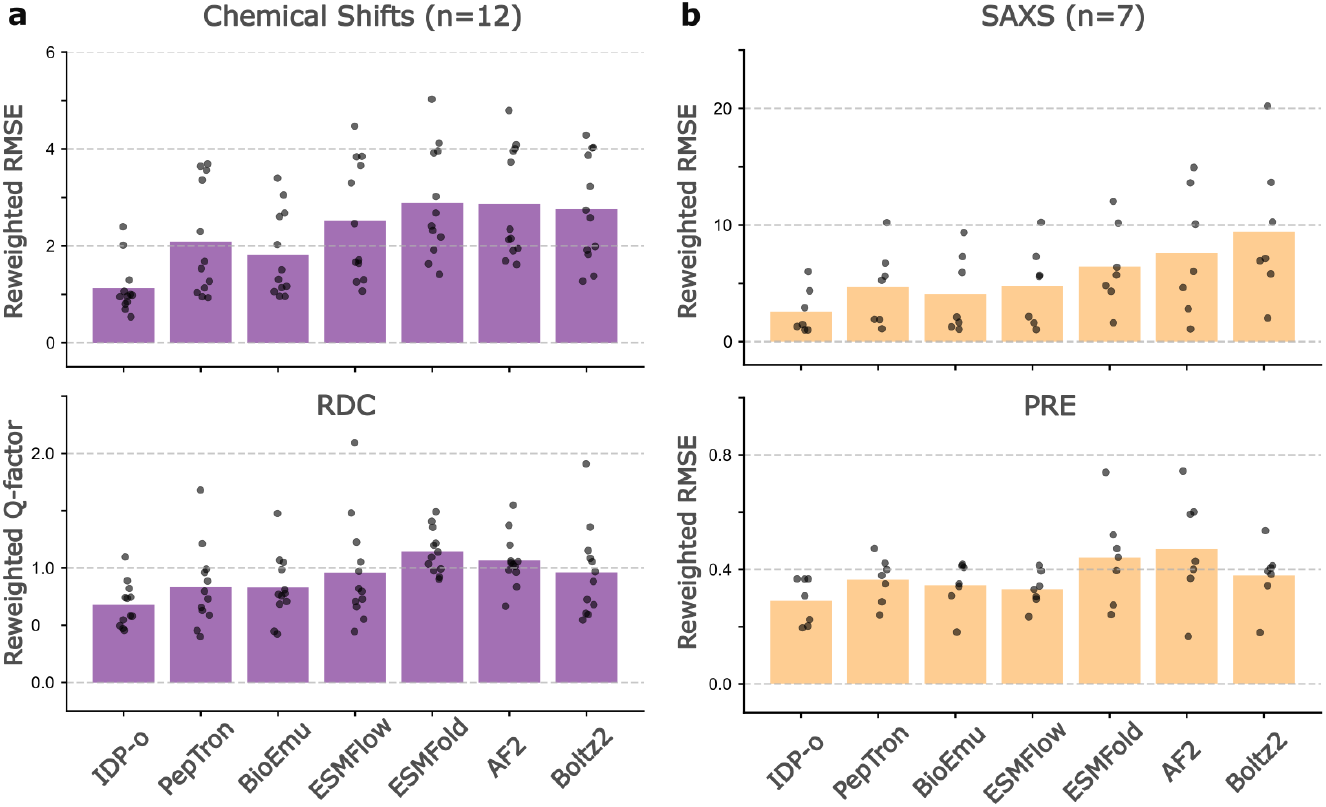
(a) RMSE between predicted and experimental chemical shifts (top panel) for ensembles generated by seven different models following maximum entropy reweighting using CS data. In the bottom panel, Q-factors measuring agreement between predicted and observed residual dipolar couplings (RDCs) across the same models and CS-reweighted ensembles are shown. Each bar represents mean performance for n=12 intrinsically disordered systems for which both CS and RDC are available; individual points correspond to results for each individual system. Lower RMSE and Q-factor values indicate improved agreement with experimental data. (b) RMSE between predicted and experimental SAXS profiles (top panel) following maximum entropy reweighting using SAXS data. In the bottom panel, RMSE of PRE data for the same models are shown. Each bar represents mean performance for n=7 intrinsically disordered systems for which both SAXS and PREs are available. Note that the RMSE values for PREs refer to the root mean squared error of predicted PRE intensity ratios (ranging from 0 to 1) relative to experimental values.

The complementary analysis using experimental data reporting on more global structural features, i.e., fitting to SAXS profiles with cross-validation restraints from PRE data (Figs. 4, D14 and D16) confirmed these trends. IDP-o again yielded the best performance, followed by BioEmu. PepTron and ESMFlow also performed well, with slightly worse agreement than that achieved by BioEmu. IdpGAN [18] delivered strong agreement for global observables (SAXS and PREs), but does not capture secondary structure preferences (Figs. D17 and D18).

Overall, similar results were obtained when comparing the models without reweighting (Fig. D14). In this case, IDP-o performed best on chemical shifts, RDCs and PREs, and second best behind IdpGAN on SAXS data. IDP-o thus appears to be particularly well suited to both capture reasonable local and global properties of disordered proteins, including in the absence of experimental reweighting. Conversely, AlphaFold2, ESMFold and Boltz2 had the worst agreement with all types of experimental data, suggesting these models do not accurately capture local or global properties of IDPs. Other ensemble-based predictors applicable to both ordered and disordered sequences typically achieve performances intermediate between those of IDP-o and AlphaFold2/ESMFold/Boltz2. The assessment of this benchmark dataset thus suggests that for typical unfolded proteins IDP-o is most suitable. Notably, RDC Q-factors achieved by IDP-o are comparable to those achieved by state-of-the-art molecular dynamics simulations of disordered proteins[26], with Q-factors ranging from ∼ 0.4 to 1.0.

Despite the overall improvements in modern ensemble predictors, this integrative benchmark also highlights persistent challenges for all models. For example, PRE experiments reveal that subtle long-range contact preferences remain difficult to predict and capture with reweighting. Residual secondary structure biases retained by some models, including for example BioEmu and PepTron, can also lead to poor agreement with experimental data (e.g., reflected in chemical shift and RDC predictions, Figs. D17 and D18). Another previously noted limitation of models such as AlphaFold2, BioEmu and PepTron is their inability to capture unfolded states of destabilized sequences that may fold into stable structures under some conditions[20]. While IDP-o captures these disordered states well, a folding bias of the other models results in large secondary chemical shifts and poor agreement with experiments (Fig. D19).

## 3 Discussion

A central contribution of this work is the explicit incorporation of disorder as a dimension in protein structure prediction. This is a critical factor widely acknowledged in the biophysics community but not yet fully integrated into current machine learning frameworks. Building on this concept, our work makes three key advances: (1) we introduce PeptoneBench, an experimentally anchored benchmark for evaluating ensemble predictions using standardized pre- and post-reweighting metrics (Fig. 1); (2) we demonstrate that fine-tuning on a disorder-focused dataset markedly improves ensemble modeling accuracy in our novel PepTron model, which utilizes a deliberately standard neural architecture (Fig. 2); and we systematically benchmark leading structure predictors, including PepTron, AlphaFold2, Boltz2, and BioEmu, with particular emphasis on performance in disordered regions (Figs. 3 and 4). Our evaluation demonstrates that the predictive accuracy of AlphaFold2 and Boltz2 systematically declines as protein disorder increases, as commonly observed. In contrast, PepTron and BioEmu sustain high performance across both ordered and intrinsically disordered protein regions. Benchmarking on ordered single-structure datasets (CAMEO22 and CASP14) highlights both Boltz2 and AlphaFold2 as leading predictors (Table 2). Collectively, our benchmark establishes robust, disorder-dependent performance baselines for the field.

Although we present a summary PeptoneBench score with and without reweighting (Table 2), we are well aware that summarizing model performance with a single score can be misleading. For this reason, we provide a range of evaluation metrics and plots as output of our benchmark, and we encourage researchers to engage with different metrics to capture distinct aspects of accuracy across datasets and levels of disorder. Moving forward, we intend to further explore the richness and nuances of the presented datasets, for example analysing how well models handle different types of IDRs, such as flexible linkers or disordered terminal domains.

While this work represents a step toward more accurate and general protein ensemble prediction, several limitations remain. First, the realism of IDR training data could be further enhanced by using either curated databases of structural ensembles [22] or long-timescale molecular dynamics (MD) simulations with improved force fields and sampling techniques [55]. Recently developed MD datasets such as ATLAS [56], mdCATH [57], DynaREPO [58], and MISATO [59] have greatly expanded the availability of simulation-based protein ensembles. However, these predominantly capture local fluctuations near the folded state and often fall short in sampling the broad conformational landscape characteristic of IDRs. This is confirmed by the fact that the ESMFlow model fine-tuned on ATLAS does not score better than the base ESMFlow model on PeptoneBench, see Fig. D11. The BioEmu MD datasets stands out by providing extensive simulations of unfolded proteins and by training explicitly on equilibrium properties [14], as reflected in its strong performance on PeptoneBench (Figs. 3 and 4).

A further challenge in ensemble validation lies in the use of forward models to translate predicted structures into experimental observables with accurate uncertainty quantification. Many such models have been optimized and validated primarily on structured systems, raising concerns about their reliability when extrapolated to disordered proteins [60, 61]. For instance, CS predictors may exhibit systematic biases in highly flexible regions, while SAXS forward models may not fully capture the effects of excluded volume in extended conformations. Comprehensive cross-validation, leveraging multimodal datasets with diverse observables, is essential to mitigate these issues; yet assembling such datasets remains difficult and resource-intensive [22, 27, 62]. Future work should prioritize the development and validation of forward models specifically calibrated for disordered protein ensembles.

Finally, it is important to acknowledge that the conformational preferences of IDPs are highly sensitive to external conditions, including temperature, pH, ionic strength, and the presence of binding partners or small molecules. Numerous studies have shown that these environmental factors can drive disorder-to-order transitions, modulate structural heterogeneity, alter phase separation behavior, and impact biological function. Achieving predictive accuracy across varying conditions is therefore critical, not just for understanding protein biophysics but also for applications in therapeutic development and drug discovery [63]. Although modeling environmental effects lies beyond the scope of this work, we highlight this challenge as a key priority for the next generation of protein structure prediction methods. Approaches that integrate physics-based modeling with learned representations, or that explicitly condition predictions on experimental variables, may prove particularly valuable in this regard.

Beyond these technical challenges, our results carry broader implications for the field. The success of relatively simple data augmentation strategies in improving disorder prediction suggests that architectural complexity and the sheer amount of training data may not be the only limiting factor. Rather, the diversity and representativeness of training data appear critical. This finding has practical implications for resource allocation in model development and highlights opportunities for community-driven efforts to curate and share high-quality ensemble datasets [64].

## Data and Code Availability

The PeptoneDB datasets, the IDRome-o dataset, the PepTron weights, and all the generated protein ensembles evaluated in the paper are available under CCBY on Zenodo at https://zenodo.org/records/17306061.

The code to reproduce the results and figures of the paper and to perform the PeptoneBench benchmark is available under Apache 2 license at https://github.com/PeptoneLtd/PeptoneBench. The code for inference and training of the PepTron model is available under Apache 2 license at https://github.com/PeptoneLtd/PepTron. The IDP-o code is available under Apache 2 licence at https://github.com/PeptoneLtd/IDP-o.

## Author Contributions

MI and CF led ideation, model development, and data analysis; JOS curated the PeptoneDB-integrative dataset, and performed analyses; FA generated the data. SB analysed data. BJ and BT contributed to research; BT and NAEV to model development, benchmarking. Additional model-development contributions from NAEV, DR, YL, and HS; All authors contributed to writing the paper.

## Acknowledgments

We thank Oliver Dutton for contributing to the ideation and implementation of IDP-o. We acknowledge Francesco Farina, Simon Chu, Farhad Ramezanghorbani, Neha Tadimeti, Emine Kucukbenli, Cedric Steenbeke, and Saee Paliwal from NVIDIA for supporting this project. We also thank the PDB, BMRB, SASBDB, UniProt and all authors of experimental studies who have publicly shared their data enabling the benchmarks presented in this manuscript.

## Appendix A PeptoneBench

### A.1 Protein Ensembles preparation

To evaluate predictions on PeptoneBench, we preprocess each protein ensemble generated by a model using a standardized protocol. Each ensemble is required to contain 100 all-atom structures. For models that do not directly output all-atom ensembles, we follow the authors’ recommended procedures where available. If guidance is lacking, cg2all [65] is used for C_*α*_-only ensembles (as with IdpGAN), and H-Packer [66] for backbone-only models (as recommended for BioEmu). No energy minimization is performed, and ensembles are stored in both pdb and xtc formats.

To ensure consistency, we apply PDBFixer https://github.com/openmm/pdbfixer to each frame, adjusting hydrogens based on the experimental pH. Each frame is saved as a separate pdb file, allowing for potential differences in protonation states across the ensemble. Subsequently, we use the appropriate forward model to calculate the experimental observable of interest for each frame, with the results recorded in a csv file.

While all forward models require individual pdb frames as input, only the original ensemble files (pdb and xtc) and the resulting csv files are stored; the intermediate, processed pdbs generated by PDBFixer are not retained.

### A.2 PeptoneDB-CS processing

Chemical shifts were sourced from BMRB via the TriZOD “moderate” selection, comprising 5,246 proteins with experimentally derived disorder annotations. The raw distribution is skewed toward ordered proteins by roughly 10:1, mirroring PDB biases. To ensure computationally tractable and balanced evaluation, we constructed a subset with equal numbers of ordered and disordered systems, yielding 659 proteins: 329 predominantly ordered and 330 predominantly disordered sequences. Disorder is quantified using a per-residue G-score [35], derived from secondary chemical shifts, where 0 indicates fully ordered and 1 fully disordered.

A delicate choice we had to make regards the forward model to compute CS from atomistic configurations. Several CS forward models have been developed, many of which exhibit comparable overall performance, with uncertainties roughly an order of magnitude larger than the experimental NMR error [43]. For our analyses, we considered both Sparta+ [42] and UCBshift [43]. (We decided not to use UCBshift-2 [67] because it did not bring any significant difference in our benchmark, since it only improves on side chains CS, and the code for that model is sill in an unmerged secondary branch of the repository.) UCBshift was selected as the default due to its greater robustness: Sparta+ occasionally produced extreme outlier predictions (sometimes exceeding 100 standard deviations relative to BMRB statistics), while UCBshift exhibited fewer such outliers and less dramatic. However, UCBshift can sometimes return *NaN* values for problematic ensemble configurations; when this occurs for any chemical shift, we discard the entire configuration from evaluation. Nevertheless, the PeptoneBench results remain qualitatively robust with respect to using UCBshift or Sparta+, as shown in Fig. A1.

It is important to note that uncertainty estimates for CS forward models are primarily derived from fully structured proteins, and uncertainties for disordered residues are typically underestimated [68]. Since experimentally resolved ensemble data for IDPs is very limited, direct quantification of uncertainties in disordered regions is challenging, and remains an unsolved problem. We take a pragmatic approach, and use as a reference point for accuracy in fully disordered regions the uncertainty estimates reported for POTENCI, a sequence-based random coil chemical shift predictor [69].

We propose a disorder-dependent uncertainty model *σ*(*a, n*) which is a function of the CS type *a*, and the residue number *n*, that interpolates linearly between the uncertainty of the forward model *σ*_*fm*_(*a*) (e.g. UCBshift) and the one of the random coil (POTENCI) *σ*_*rc*_(*a*), using the per-residue g-score *g*(*n*):

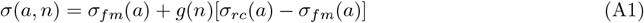

This uncertainty matches the reported forward model uncertainty *σ*_*fm*_(*a*) for residues belonging to a structured region, and the smaller one of the POTENCI random coil predictor for IDRs. Our uncertainty model is validated by the RMSE results, see Sec. D, which approach one sigma for the best models. If one is to naively use *σ*_*fm*_(*a*) instead of our *σ*(*a, n*) to compute the RMSE, the resulting curves would be way below one most of the time for disordered regions, even for poor protein prediction models, a clear sign of underestimated uncertainty. We stress that the reported RMSEs are mostly useful for comparisons between models, and their absolute value should not be trusted blindly, since they are dependent on the chosen uncertainty model.

**Fig. A1.**
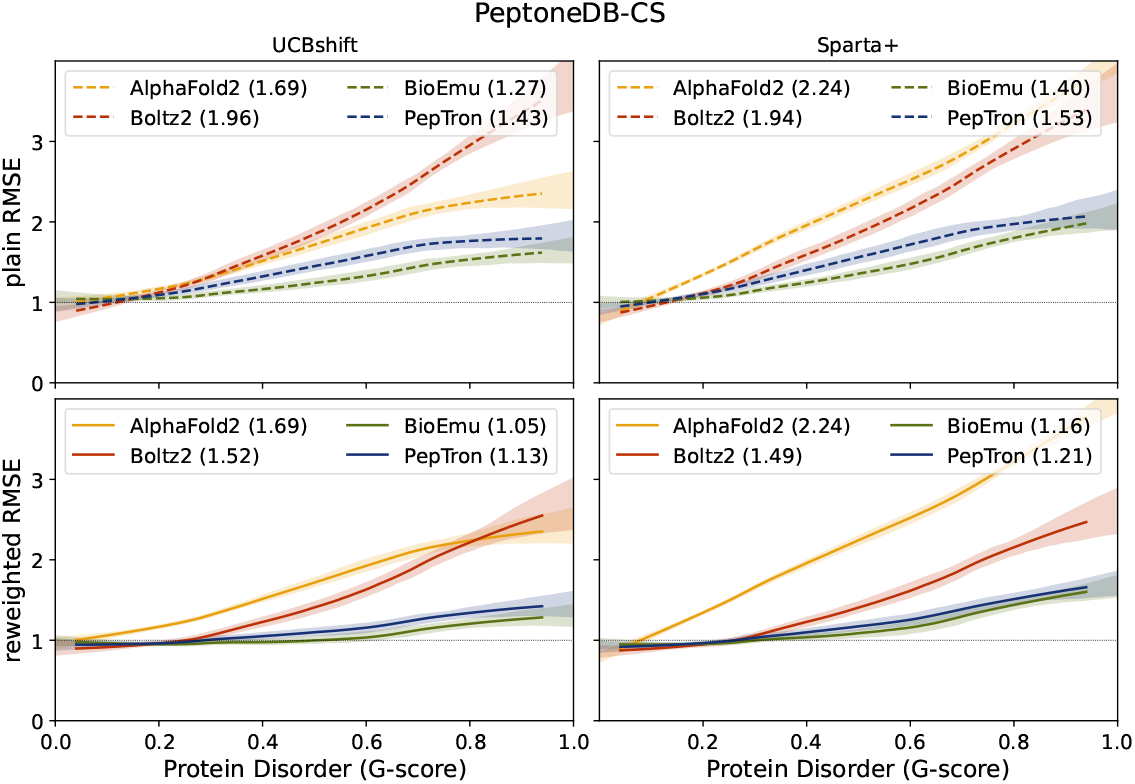
Results on the chemical shift database for some of the considered generative models, using two different forward models, UCBshift [43] and Sparta+ [42]. The values in the parenthesis are the area under the LOWESS curve, as in Fig. 3.

### A.3 PeptoneDB-SAXS processing

Entries were curated from SASBDB [39], retaining only monomeric proteins composed exclusively of standard amino acids, with sequence lengths between 10 and 500 residues, and measurements conducted at pH 4.0–10.0. A python notebook to reproduce this filtering is provided in the accompanying repository. Instead of processing raw SAXS data directly, we used the SASBDB-supplied preprocessed files (generated with GNOM [70]), ensuring consistent *q*-range filtering and omitting any entries lacking these files. Contrary to the chemical shifts case, uncertainties from experiments are non-negligible, so we use BIFT [71] to rescale them, to reduce the variance in how different groups report them. The final dataset comprises 439 sequences, with no adjustment for disorder content. Redundancy was minimized by clustering with MMseqs2 [72] (`easy-cluster` mode; 90% sequence identity threshold), removing highly similar sequences.

For reweighting, we generally follow the procedures outlined in Ref. [73], with minor modifications. Scattering profiles are computed from atomistic structures using Pepsi-SAXS [44] as the forward model. As with the NMR chemical shift calculations, PDBFixer (https://github.com/openmm/pdbfixer) is used to add hydrogens to each frame according to the experimental pH. Pepsi-SAXS is then run with default parameters on each frame; unlike Ref. [73], we allow the fitting of *r*_0_ and *δρ*_0_ to vary for every ensemble frame. While these parameters may indeed depend on protein conformation, fitting them individually introduces a risk of overfitting and can yield physically inconsistent values across frames. Ideally, *r*_0_ and *δρ*_0_ should be estimated using a physical prior, though this is left for future work. Following the approach advocated in the Pepsi-SAXS paper, we prioritize achieving the best possible agreement with experimental data, even at the expense of potential overfitting. The default option of *cst* = 0 in Pepsi-SAXS is maintained, and we do not apply corrections for overall shifts in the SAXS profile.

Since scattering intensity is reported in arbitrary units, we first compute a weighted average of the intensities over all ensemble frames and then rescale the result to match the experimental profile using the formula described by Svergun (1995):

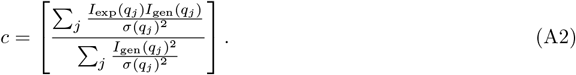

And then compute the *χ*^2^ as

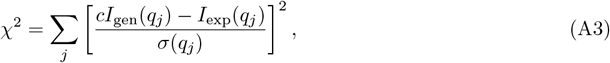

This introduces a further dependence on *I*_gen_(*q*) that breaks the relation between **ℒ**_*θ*_(**w**) and Γ_*θ*_(***λ***), so in this case we run the minimization in the space of the weights **w**, rather than in ***λ*** as in Ref. [28] (see Sec. A.5).

**Table A1.** Disordered protein datasets used in the PeptoneDB-Integrative benchmark.

| Protein | Datasets | # of residues | Average G-score |
| --- | --- | --- | --- |
| Aβ40 | CS[74], RDC[75] | 40 | 0.784 |
| ACTR | CS[76], RDC[76], SAXS[77], 4x PRE[76] | 71 | 0.770 |
| ANAC046 | CS[78], SAXS[79], 3x PRE[80] | 167 | 0.814 |
| αSyn | CS[81], RDC[82], SAXS[83], 5x PRE[84] | 140 | 0.841 |
| Δ131Δ | CS[85], RDC[86] | 131 | 0.674 |
| drkN | CS[87], RDC[88], SAXS[89], 2x PRE[90] | 59 | 0.597 |
| ERNTD | CS[91], SAXS[91], 6x PRE[91] | 184 | 0.802 |
| FLN5A3A3 | CS[92], RDC[92], SAXS[92], 6x PRE[92] | 114 | 0.868 |
| NTail | CS[93], RDC[94] | 132 | 0.802 |
| OPN | CS[95], RDC[96] | 220 | 0.848 |
| p15PAF | CS[97], RDC[97], SAXS[97] | 110 | 0.879 |
| p53NTD | CS[98], RDC[99] | 102 | 0.770 |
| PaaA2 | CS[45], RDC[45], SAXS[45] | 71 | 0.432 |
| RCFTR | CS[100], RDC[88] | 189 | 0.772 |
| Sic1 | CS[101], RDC[101], SAXS[102], 6x PRE[101] | 92 | 0.869 |
| Tau | CS[103], RDC[104], SAXS[105], 10x PRE[104] | 441 | 0.859 |

### A.4 PeptoneDB-Integrative processing

We curated from the literature a set of 16 proteins (see Table A1) that either have matched NMR chemical shifts and ^1^*D*_HN_ RDCs or SAXS and PREs, extending prior integrative benchmarks[15, 26, 40]. The matched modalities enable cross-validation of local and global features, respectively, differentiation between calibratable errors (addressable by reweighting) and sampling deficiencies (lack of requisite conformations), and quantitative assessments of maximum-entropy reweighting behaviour on realistic IDP systems. For all proteins in this dataset, we used ensembles of 1,000 conformations. NMR chemical shifts and SAXS data were calculated and compared with experimental values as described in the previous sections.

RDCs were calculated using PALES[106] and a local alignment window of 15 residues[88]. After calculating a weighted ensemble average for each ensemble, the magnitude of the predicted RDCs was scaled to optimise the agreement with experimental data, using only residues whose predicted RDCs matched the sign of the experimental values[20]. The agreement with experimental data was quantified using the RDC quality factor (Q-factor):

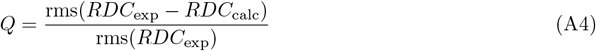

PREs were calculated using the MTSL spin label rotamer library implemented in DEER-PREdict[46]. Agreement between calculated and experimental data was quantified using a regular RMSE using intensity ratios 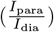. To account for the uncertainty of the nucleus-electron vector correlation time, *τ*_*C*_, this parameter was optimised against the experimental data by scanning values in a range of 1-20 ns in steps of 1 ns. Calculated (and if available experimental) PRE rates, Γ_2_, were converted to intensity ratios using a transverse relaxation time, R_2_, of 10 s^−1^ and INEPT time of 10 ms. For one dataset (FLN5A3A3) where PRE intensity ratios were measured using a HMQC rather than a HSQC experiment[92], we used the following equation to convert Γ_2_ to intensity ratios:

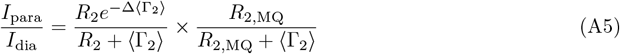

Here, R_2_ and R_2,*MQ*_ (the linewidth contribution from multiple-quantum relaxation in the indirect dimension) are both set to 50 s^−1^ and the delay (Δ) to 10.86 ms.

For reweighting with NMR chemical shifts, we used 64 values of *θ* and to avoid overfitting we employed cross-validation restraints from RDC experiments. The optimised posterior ensembles were chosen based on having optimal agreement with the orthogonal RDC data, while satisfying an effective sample size of at least 100 conformers (10% relative effective sample size). If the agreement with the orthogonal data decreased for all posterior ensembles, the prior was used. The same principle was applied to ensemble reweighting with SAXS data, where PREs were used as cross-validation restraints. Finally, to fairly compare all ensemble prediction models we calculated the average metrics for each experimental data type across all proteins that were successfully predicted by all models.

### A.5 Reweighting to a target effective sampling size

Following the implementation of Ref. [31], we define the root mean squared error (RMSE) for a set of *M* experimental observables 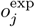 with uncertainty *σ*_*j*_ and an ensemble of *N* protein samples **x**_*i*_ with weights *w*_*i*_ and forward model outputs 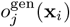 as:

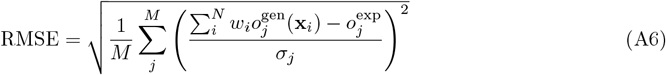

with 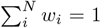. We then optimise the objective function that balances adherence to the experimental data with maximum entropy weights via the parameter *θ*

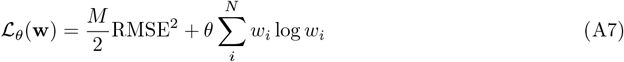

with respect to the weights *w*_*i*_.

Alternatively, optimization can be performed with respect to the Lagrange multipliers *λ*_*j*_ of

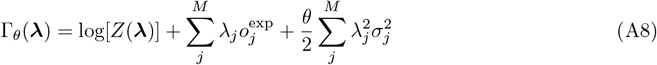

where

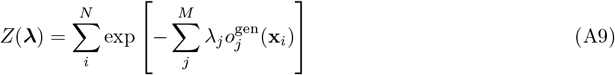

and

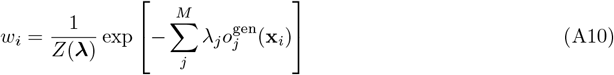

and assuming Gaussian, independent errors *σ*_*j*_ which include both experimental and forward model contributions.

The regularization parameter *θ* controls the tradeoff between fitting the data and maintaining ensemble diversity: smaller *θ* enforces a tighter fit at the risk of overfitting.

A useful metric for monitoring diversity is the Kish effective sample size (ESS):

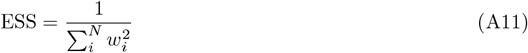

which reports the effective number of frames contributing to the ensemble average. A typical RMSE versus *θ* curve is shown in Fig. A2. Ideally, the regularization parameter *θ* is selected to prevent overfitting, for example by using cross-validation with independent datasets.

For the PeptoneDB-CS and PeptoneDB-SAXS datasets, independent sets of experimental data are not available, preventing a direct test for overfitting through cross-correlation. Our primary objective in the context of the present work is to assess whether an ensemble is “reweightable”. We therefore set a threshold on the ESS to avoid solutions where only a small number of frames dominate the ensemble. We set the target ESS to 10% of the total number of frames, and optimize *θ* over a grid, refining the value such that ESS 0.1≈ *N* using a root-finding algorithm.

A relevant issue not encountered in molecular dynamics simulations is that machine learning–based protein ensemble generators can produce unphysical samples, such as broken chains or severe atomic clashes. The forward models used to compute experimental observables (e.g., chemical shifts or scattering profiles) typically do not screen for such artifacts. Consequently, it is necessary to introduce a filtering, in which any frame displaying unphysical features is excluded by assigning it zero weight.

For this purpose, we apply the criteria used by BioEmu: a maximum distance of 0.45 nm between contiguous C-*α* atoms, a maximum of 0.2 nm between contiguous backbone carbon and nitrogen atoms, and a minimum distance of 0.1 nm between any two atoms. While such unphysical frames are generally rare, their occurrence can be significant for certain generators or specific proteins, sometimes resulting in more than 90% of frames being filtered and effectively precluding reweighting refinement. Although the presence of these artifacts is not problematic by itself, their impact is to reduce the generator’s effective sampling efficiency and, consequently, its performance on the benchmark.

**Fig. A2.**
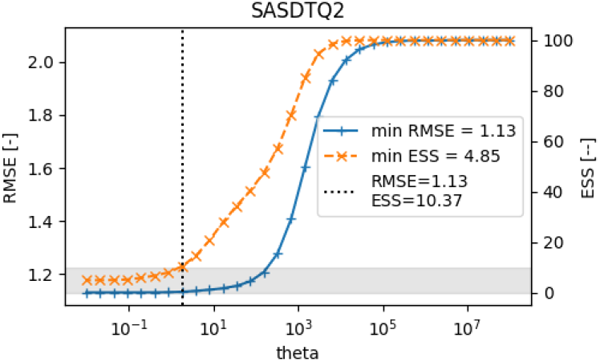
Example of RMSE (blue, left axis) and effective sample size (ESS, orange, right axis) as functions of the regularization parameter *θ* during ensemble reweighting for an example SAXS entry. The minimum RMSE and ESS values are highlighted, and a vertical dotted line indicates the chosen *θ* corresponding to an effective sample size of approximately 10% of the ensemble. This plot illustrates the tradeoff between fit to the experimental data (low RMSE) and ensemble diversity (high ESS) as *θ* varies.

## Appendix B IDP-o

To efficiently generate physically plausible ensembles of intrinsically disordered proteins (IDPs) at scale with minimal computational cost, we developed IDP-o, a fragment-based ensemble generator. Like previous fragment-based approaches [47, 48], IDP-o constructs protein ensembles by assembling fragments from a structural database. However, rather than relying on the limited structural diversity of the PDB, IDP-o leverages the comprehensive AFDB-UniProt database, which encompasses over 214 million AlphaFold-predicted protein structures [9]. This vast repository enables the consistent use of 6-residue fragments, ensuring more faithful representation of local secondary structure elements while maintaining computational efficiency.

To enable practical large-scale use, AFDB-UniProt is compressed using Foldcomp [107] and reconstructed with NeRFax [108], an efficient approach for transforming internal coordinates into Cartesian space. This strategy reduces the database footprint to approximately 1.1 TB while preserving rapid query performance, allowing efficient retrieval of structural ensembles for peptide fragments.

The IDP-o workflow consists of three core components (Fig. B3):

1. **Fragment-Based Ensemble Query System:** Input protein sequences are partitioned into overlapping 6-residue fragments, which are used to query the compressed structural database to identify natural conformational matches.
2. **Structure Clustering and Alignment:** Retrieved fragment structures are clustered based on C_*α*_ torsion angles to group conformationally similar hits. Each cluster is represented by a centroid conformation, with stochastic overlays providing discrete conformational states for individual fragments.
3. **Ensemble Assembly from Multiple Fragments:** Full-length ensembles are reconstructed by combining structural clusters across sequential fragments. Because each fragment typically provides multiple structural alternatives, combinatorial assembly generates diverse global conformations. While local structural coherence is maintained through alignment constraints, global flexibility and heterogeneity are preserved.

On the PeptoneBench, IDP-o is the best IDP generative model among the ones we tested, as shown in Fig. D9.

### B.1 IDRome-o dataset

The IDRome-o dataset was created using IDP-o to produce protein ensembles (100 conformations each) for sequences extracted from the IDRome [34] sequences database. Only entries between 30 and 300 residues in length were considered. These were further clustered in sequence space using MMseqs2 [72] default `easy-cluster` parameters. The final structural database assembled using IDP-o comprises 100 samples for each of 22, 399 IDR sequences, for a total of 2, 060, 934 residues. The creation of IDRome-o took approximately 30 days on a A100 DGX box (8 GPUs, 128 CPUs). In Fig. B4 we report the average gyration radius for the IDRome dataset as estimated from IDP-o (no reweighting) and CALVADOS coarse grained MD [34].

**Fig. B3.**
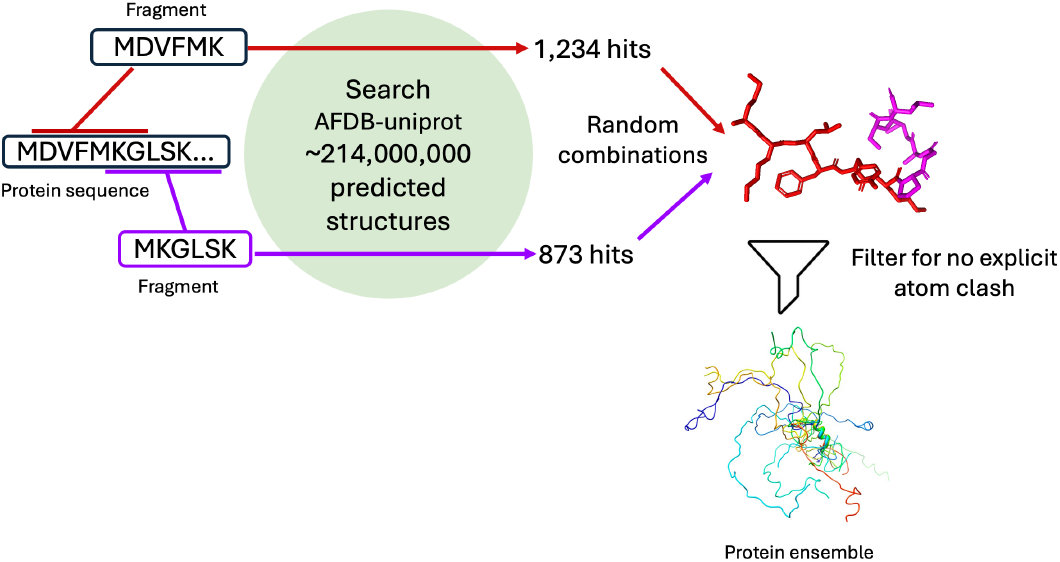
IDP-o fragment-based ensemble generation workflow. The protein sequence is decomposed into overlapping 6-residue fragments. Each fragment is queried against the AFDB-Uniprot (214M) database yielding thousands of structural hits. Random combinations of fragment conformations are assembled and filtered to remove structures with explicit atom clashes, generating diverse protein ensembles that maintain realistic local stereochemistry while exploring the full conformational space accessible to intrinsically disordered proteins.

**Fig. B4.**
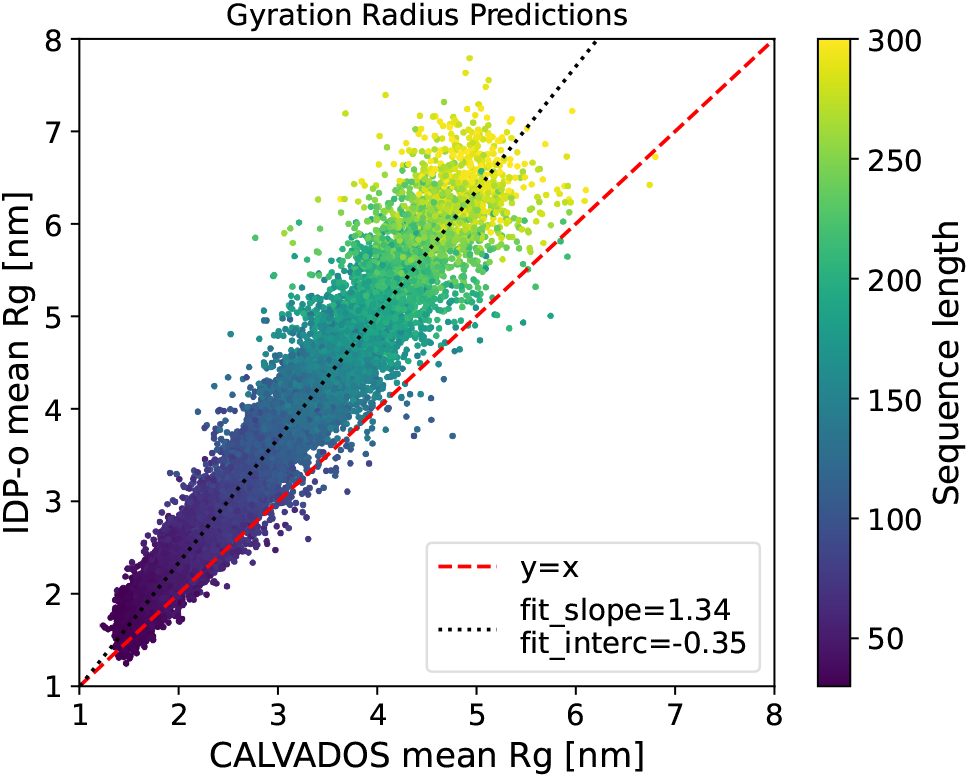
Average radius of gyration (Rg) for IDRome ensembles. Ensembles were generated using IDP-o and CALVADOS coarse-grained MD [34]. IDP-o, consistent with other fragment-based methods, shows a tendency to sample more extended structures, particularly for longer sequences.

## Appendix C PepTron

Our approach builds upon the flow matching framework for protein structure generation introduced by Jing et al. [13], adapting their ESMFlow architecture for end-to-end training on multidomain protein ensembles. The model architecture follows the core design principles established in AlphaFlow, which repurposes pre-trained single-state structure predictors by integrating continuous flow matching to generate conformational ensembles. We call this model PepTron.

The architecture comprises several key components, described below.

1. **Sequence Encoding**: input amino acid sequences are embedded using the pre-trained BioNeMo `ESM2-3B`^1^ transformer model by NVIDIA which shows improved performance over the pure-pytorch baseline [49, 109]. This model comprises 36 transformer layers with approximately 3 billion parameters, and was trained on UniProt 2021 04 sequence database[110], following the protocol described in the BioNeMo framework^2^. The resulting contextualized sequence representations are combined via a learned weighted sum across all layers and further augmented with trainable amino acid embeddings [51].
2. **Folding Trunk:** The core processing occurs through 48 triangular self-attention blocks that iteratively refine both sequence (*s*_*s*_) and pairwise (*s*_*z*_) representations through cross-attention mechanisms and triangular multiplicative updates[51]. In order to tackle biological systems made up of thousands of residues, we integrated the triangular attention cuEquivariance kernel^3^ which provides memory complexity reduction from *O*(*N* ^3^) to *O*(*N* ^2^) with *N* number of residues.
3. **Structure Module:** The refined representations are passed to an invariant point attention (IPA)-based structure[111] module that predicts backbone frames, side-chain conformations, and atomic coordinates.
4. **Time-Conditioned Pairwise Processing:** Pseudo-*β* coordinates (C*β* atoms or C*α* for glycine) are used to compute pairwise distance matrices, which are binned into distograms and processed through input pair stacks. Crucially, a continuous time variable *t* is embedded using Gaussian Fourier projections and added to the pairwise representation, enabling the model to condition on different stages of the denoising process, as described in [13].

A key advantage of this model compared to previous approaches is its exceptional usability at inference time. Unlike models that require precalculated embeddings, Multi-Sequence-Alignments (MSAs), or downloading separate ESM2 or ESMFold weights, this architecture allows prediction using only a single, freely available model checkpoint. Users simply provide a list of sequences to generate ensembles, with no need for additional preprocessing steps or resources. The PepTron codebase with the related check-point also supports resuming training if desired, making the workflow highly flexible and accessible. This design effectively makes protein ensemble prediction straightforward and accessible.

A key characteristic of PepTron is its ability of modeling proteins at any point of the order-disorder continuum, as shown in the half-disordered protein in Fig. C5.

### C.1 Training

PDB training data are sourced from a snapshot taken in May 2020, with sequences clustered at 40% identity using MMSeqs2 [72]. Training is performed using random crops of length 512 with a batch size of 64, and omitting both recycling and the use of templates. PepTron is optimized on a combination of FAPE, pLDDT, distogram, and supervised *χ* angle losses. To ensure accuracy in initial structure prediction, half of all training samples are provided with clean inputs *t* = 0, while the remaining half are given noisy initializations. Additionally, half of the samples use predictions from earlier iterations as additional input (self-conditioning), improving stability and accuracy. Training performance is tracked with both precision and conformational diversity metrics on a validation set consisting of 183 CAMEO[113] targets deposited between August and October 2022, following the protocol in Jing et al. [114].

For fine-tuning on IDP-o ensemble data (IDRome-o), optimization resumes from the best PDB-trained checkpoint. The training set is made up of two different datasets, IDRome-o, which accounts for the 70% of it and the PDB dataset described above, comprising the remaining 30%. To avoid catastrophic forgetting and ensure the fine-tuned model could properly learn representations along the order-disorder continuum, we deliberately decided to train the model on both structured and disordered proteins, as our ultimate goal was to cover the full spectrum from order to disorder. The effect of the fine-tuning can be seen in Fig. C6.

The training configuration is otherwise unchanged except that targets are sampled uniformly (each with a random conformation), the proportion of clean initialization *t* = 0 is reduced to 10% and the self-conditioning feature is disabled entirely, so the model does not use previous predictions and instead relies only on the current input and target. This adjustment is intended to maximize diversity and ensure unbiased ensemble generation for intrinsically disordered proteins. During this phase, progress is monitored by the loss on the validation split.

### C.2 Inference

The structures generated with PepTron for this work have been obtained at ten uniformly spaced denoising steps from *t* = 0 to *t* = 1, always beginning with a clean, noise-free input for the first prediction. Structure generation for each member of the predicted ensemble is parallelized across available GPUs, maximizing GPU memory occupancy and delivering average speedup of about 10x compared to non-parallelized implementations. This design ensures fast, efficient and most importantly scalable ensemble prediction even for large ensembles. In addition, the flow matching framework has been modularized through the integration of NVIDIA MoCo^4^, which makes it a plug-and-play block, allowing users to easily substitute or benchmark different flow matching and diffusion methods with minimal code changes, thus enhancing flexibility and extensibility in generative protein modeling.

**Fig. C5.**
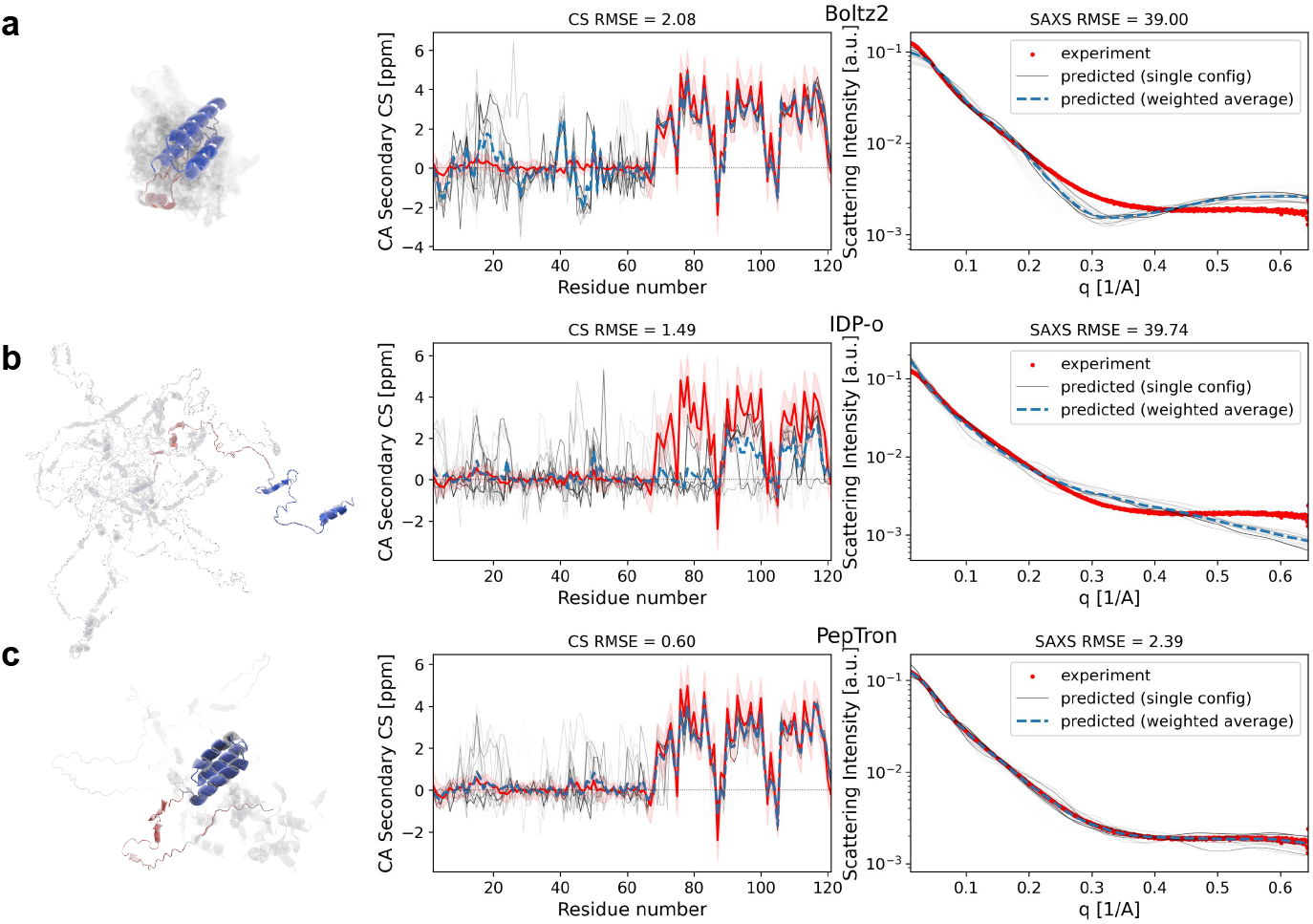
Comparative structural predictions and experimental validation for Menangle virus phosphoprotein [112]. Structural predictions for a partially unstructured protein (mean G-score = 0.46) are compared against experimental NMR CS and SAXS data. (a) Boltz2 prediction generates structures with high helical content, evident in both the 3D model and elevated secondary chemical shift values (CS RMSE = 2.08), resulting in poor agreement with experimental SAXS profiles even after reweighting (SAXS RMSE = 39.00). (b) IDP-o prediction produces extended conformations that lack the three helical structures in the C-terminal domain (CS RMSE = 1.11), showing partial chemical shift agreement and poor SAXS fitting (SAXS RMSE = 2.39). (c) PepTron prediction captures the partially unstructured character of the protein, demonstrating better agreement with both experimental datasets (CS RMSE = 0.60, SAXS RMSE = 2.39) and providing the most realistic structural ensemble.

### C.3 Computational cost

Experiments were conducted on a system provided with 8x NVIDIA H200 GPUs and 128x Intel Xeon Platinum 8468 logical cores. The total training cost summarized in Table C2.

**Table C2.** PepTron training cost.

| Model | Dataset | # Samples | Training Time (h) | Avg. Step Time (s) |
| --- | --- | --- | --- | --- |
| PepTron-base | PDB | 0.8M | 157 | 37 |
| PepTron | PDB | 0.2M | 185 | 38 |
|  | IDRome-o | 0.6M |  |  |

A critical aspect of our model is its computational efficiency, particularly for generating large structural ensembles. We analysed the inference time of our model with and without the NVIDIA cuEquivariance-optimized kernels. As illustrated in Figure C7, the PepTron cuEquivariance model dramatically speeds up inference time compared to the baseline PepTron implementation. We tested the following proteins (UniProt ID): P0CX86, P00974, P0CG47, P04637, and Q6ZU69; choice driven by their structural diversity and size. This performance gain is primarily due to a fundamental improvement in computational scaling. The practical implications of this acceleration are most evident in our novel inference protocol, where ensemble generation is parallelized on GPUs. The enhanced scaling makes it feasible to generate hundreds of ensemble structures in parallel for small-to medium-sized proteins (up to ∼ 200 residues), and tens of structures for larger proteins, all parallelised on the available devices.

**Fig. C6.**
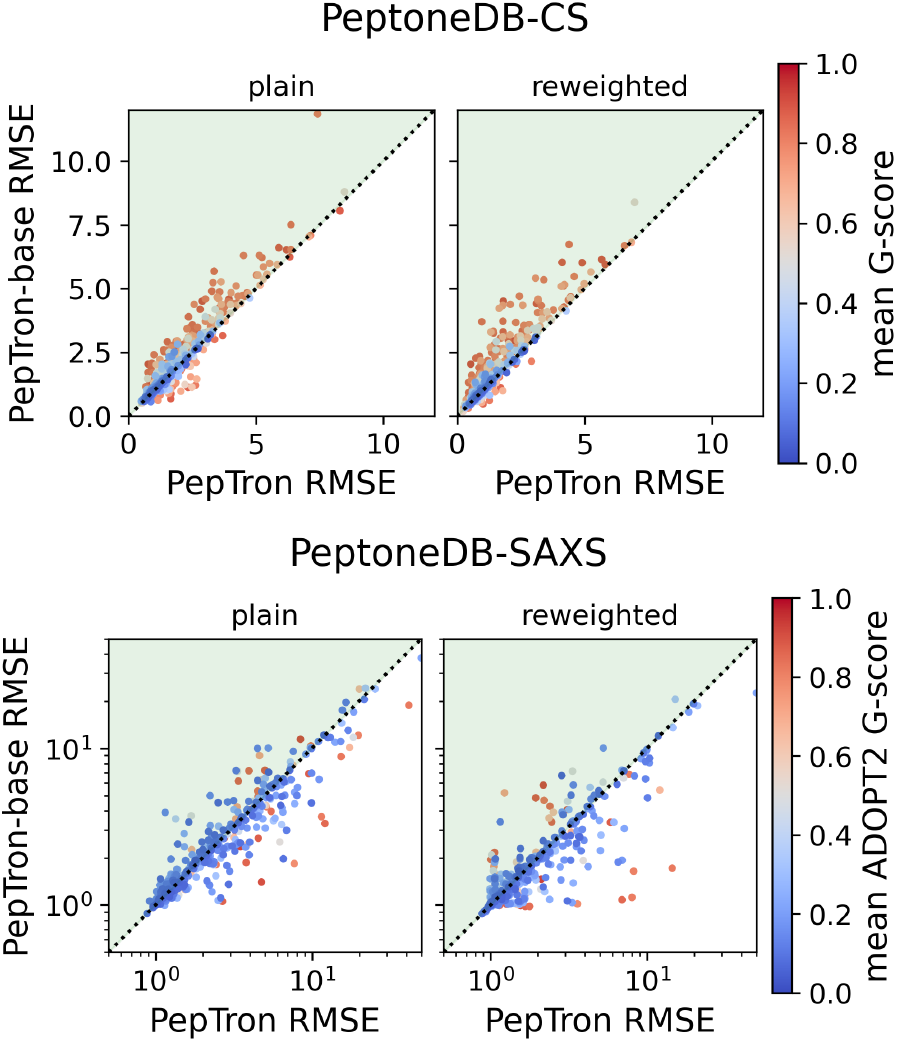
The effect of fine-tuning the PepTron model with IDRome-o. Each point shows the average RMSE (with or without reweighting) of a predicted protein ensemble with respect to the experimental data in the PeptoneDB. Ensembles in the upper left part of the plots (highlighted in green) have improved with the fine-tuning.

**Fig. C7.**
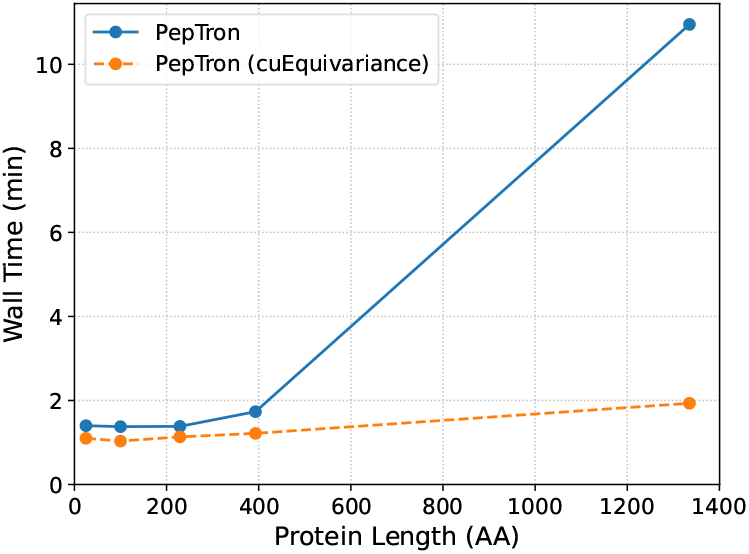
PepTron inference cost for a single protein structure. Inference time in minutes as a function of protein amino acid sequence length for the baseline PepTron model (solid line) and the version with cuEquivariance attention kernel integrated (dashed line). The cuEquivariant implementation shows substantially improved scaling, reflecting the reduction in computational complexity from cubic to quadratic.

## Appendix D Benchmarking protein generative models

We select a total of 12 popular machine learning generative model that generate an ensemble of protein 3D conformations from the protein sequence, and test them on PeptoneBench. Each model output is processed as explained in Sec. A.1. Except for the first two models, we generated 100 frames ensembels for each entry of the PeptoneDB-CS and PeptoneDB-SAXS, and 1000 frames for PeptoneDB-Integrative dataset.

For each model we compute the LOWESS regression, as implemented in `statsmodels`^5^ library, with parameters `it=1` and `frac=0.5`, and estimate the uncertainty at 95% confidence interval with 200 bootstrapping resamples.

Figure D8 reports two models that are not intended to generate an ensemble, but rather a single structure. In this special case of an ensemble of one, reweighting has no effect.

- **AlphaFold2** [9]. We use the ColabFold batch implementation [115] and run with default parameters, taking the best of 5 predictions.
- **ESMFold** [116]^6^. We ran ESMFold with default parameters.

Figure D9 presents four models which are specifically developed for IDP ensemble generation.

- **IDP-o** Knowledge-based model introduced in this work, see Sec. B.
- **IdpGAN** [117]^7^. Coarse-grained based model, cg2all [65]^8^ was used to obtain all-atom ensembles. We ran IdpGAN with batch size = 16
- **IdpSAM** [118]^9^. Coarse-grained based model, cg2all [65]^10^ was used to obtain all-atom ensembles.
- **IDPFold** [119]^11^. Backbone-only model, H-Packer [66] was used to obtain all-atom ensembles through the `bioemu.sidechain_relax` script provided by BioEmu^12^, using options `--no-md-equil --md-protocol md_equil`.

Four other models natively support protein ensemble generation.

- **ESMFlow**^13^ [13]. The PDB version^14^ with default inference parameters.
- **ESMFlow-MD**^15^ [13]. The version fine-tuned on the ATLAS MD dataset^16^, with default parameters.
- **BioEmu** [14]^17^. Backbone-only model, H-Packer [66] was used to obtain all-atom ensembles. We ran BioEmu with `--filter_samples=False` and a self-hosted MSA server^18^. We used the `bioemu.sidechain_relax` with options `--no-md-equil --md-protocol md_equil` for sidechain reconstruction.
- **Boltz1x** [120]^19^. We ran Boltz1x with `--output_format pdb` and a self-hosted MSA server^20^
- **Boltz2** [53]^21^. We ran Boltz2 with `--output_format pdb` and a self-hosted MSA server^22^

Finally we considered PepTron before and after the fine-tuning on IDRome-o, see Sec. C.

- **PepTron-base** [our]. The model trained only on the PDB.
- **PepTron** [our]. The final model, after fine-tuning.

### Note on predictor failures

In certain relatively rare cases, we have not been able to successfully generate predictions for certain model /protein combinations. More specifically:

- **BioEmu**: sidechain reconstruction failed for a total of 1 entry in the CS dataset and 3 entries in the SAXS dataset.
- **Boltz2**: model went out of GPU memory when trying to generate 100 frames ensembles for a total of 2 entries in the CS dataset and when generating 1000-frames ensembles for 3 entries in the Integrative dataset, on a H200 GPU with 141GB of VRAM.
- **IDP-o**: model fails if one of the 6-residues-long fragments is not present in the reference database. This is rare enough that we have not implemented a backup strategy like moving to shorter fragments.

**Fig. D8.**
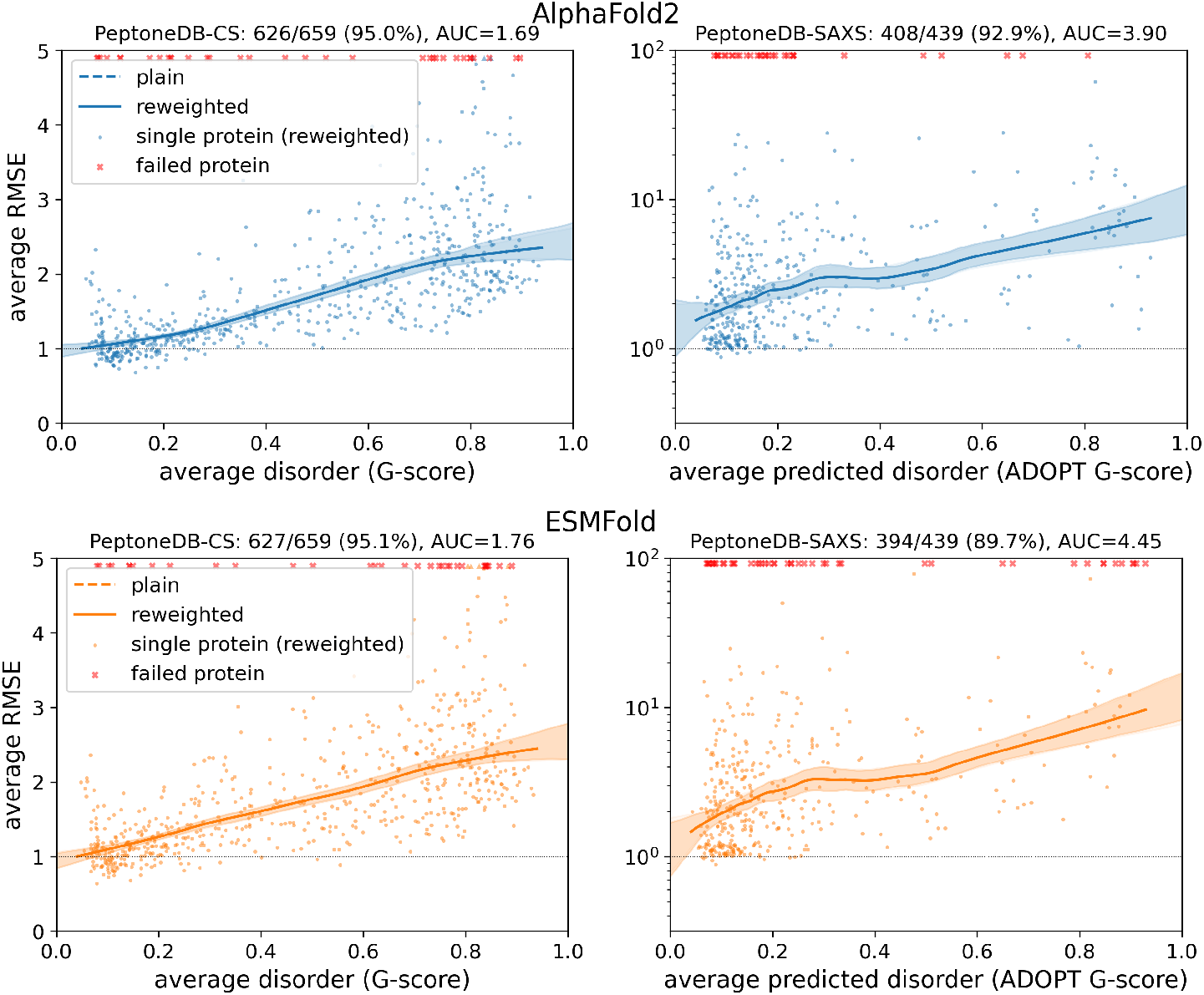
PeptoneBench results for AlphaFold2 and ESMFold. These models only output one single structure per protein, so reweighting has no effect on the results.

In other cases the models does generate ensembles, but none of the generated frames passed the physicality filter, meaning that either the protein chain is broken or some atoms are overlapping. These entries are also regarded as failed, since no configuration is usable to compare with experiments.

For convenience, we introduce a PeptoneBench summary score *s* (the lower the better):

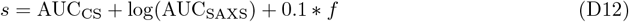

where *f* is the fraction of failed entries out of the total, averaged over the CS and SAXS datasets. An entry is considered failed if the proteins ensemble generation was not successful or if it did not contain any sample that passes the physical filtering step (due to broken backbone or atom clashes). The AUC could be either from the plain or the reweighted LOWESS curve. The PeptoneBench score would be equal to 1 or smaller for a model that generates ensembles completely in agreement with the experimental data. PeptoneBench scores for all the models evaluated are reported in Table D3 and Fig. D13.

### D.1 Results on the integrative dataset

We have assessed 9 generative models before and after reweighting on the PeptoneDB-Integrative dataset. The results and ranked models for all experimental parameters (chemical shifts, RDCs, SAXS, PREs) are shown in Fig. D14.

**Fig. D9.**
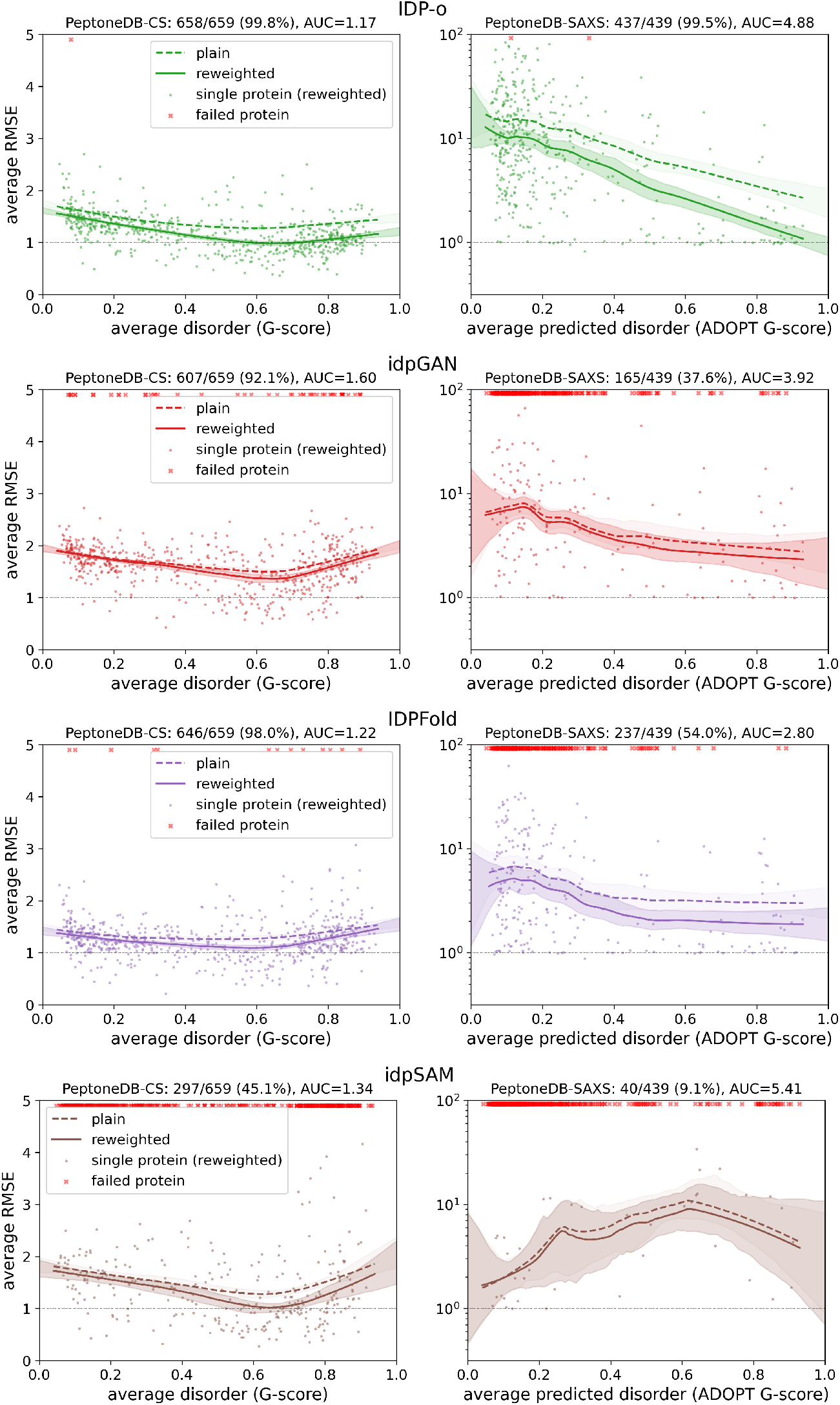
PeptoneBench results for IDP-focused generative models. These models tend to struggle to produce physically valid configurations on the long structured proteins of the PeptoneDB-SAXS, in particular IDPsam, which was trained only on short protein sequences. IDP-o does not fail there, but produces configurations that are too extended, with poor compatibility with SAXS data.

**Fig. D10.**
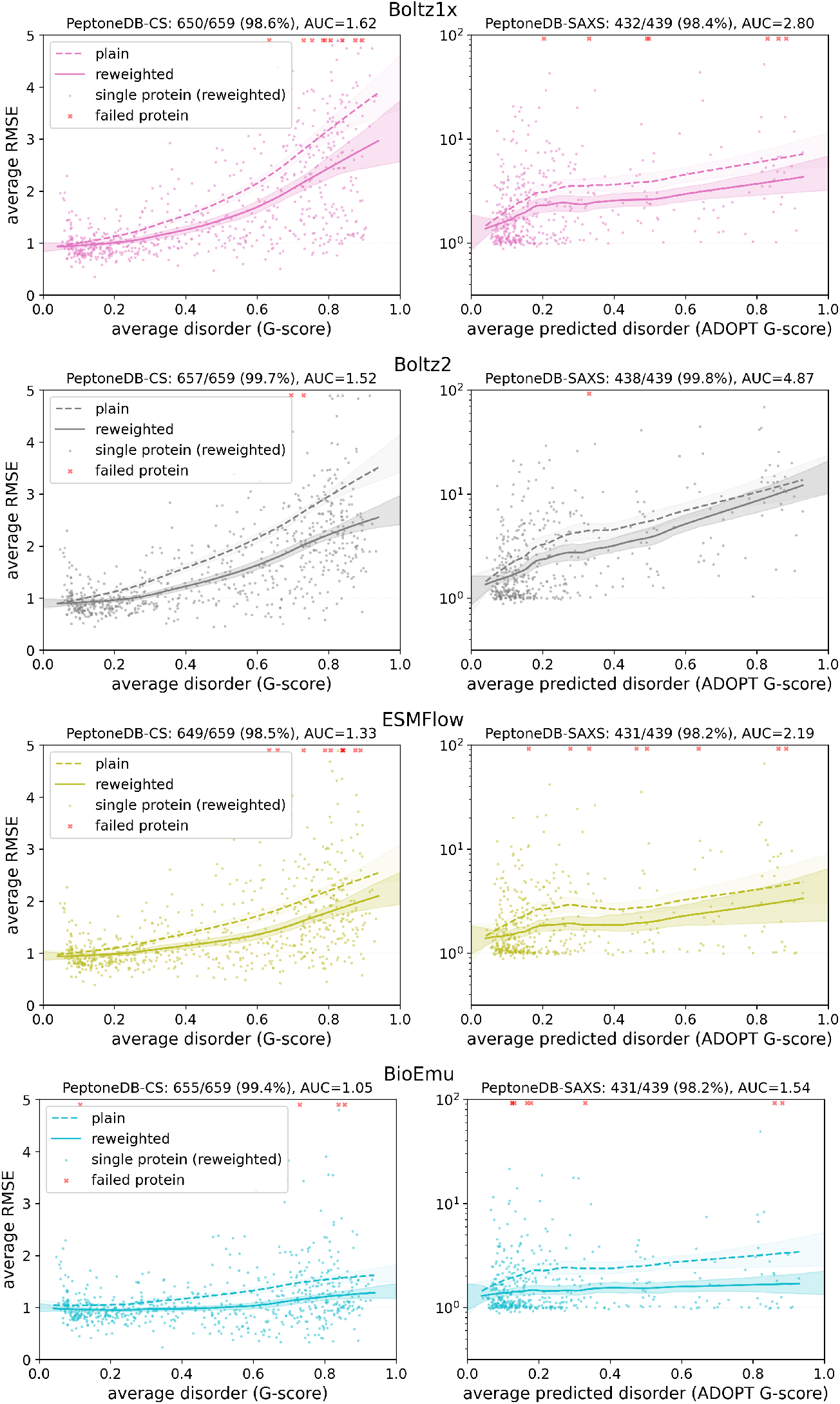
PeptoneBench results for some of the most popular protein ensemble generative models. It is interesting to notice that Boltz2 improves on the CS dataset, over Boltz1x, but gets worse on the SAXS one. BioEmu is currently the best overall model.

**Fig. D11.**
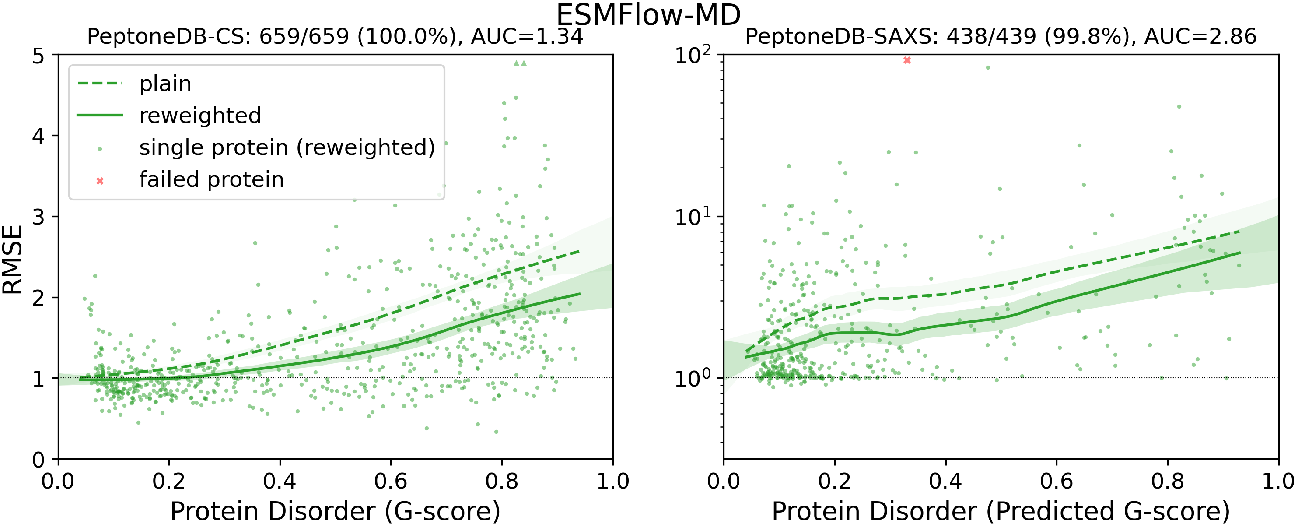
PeptoneBench results for ESMFlow fine-tuned on the ATLAS MD dataset [56], presented in Ref. [13]. Contrary to IDRome-o, the ATLAS dataset is not rich in IDRs, and using it for fine-tuning does not improve performance on PeptoneBench. In fact, the resulting model has a slightly worse score, see Table D3 and Fig. D13.

**Fig. D12.**
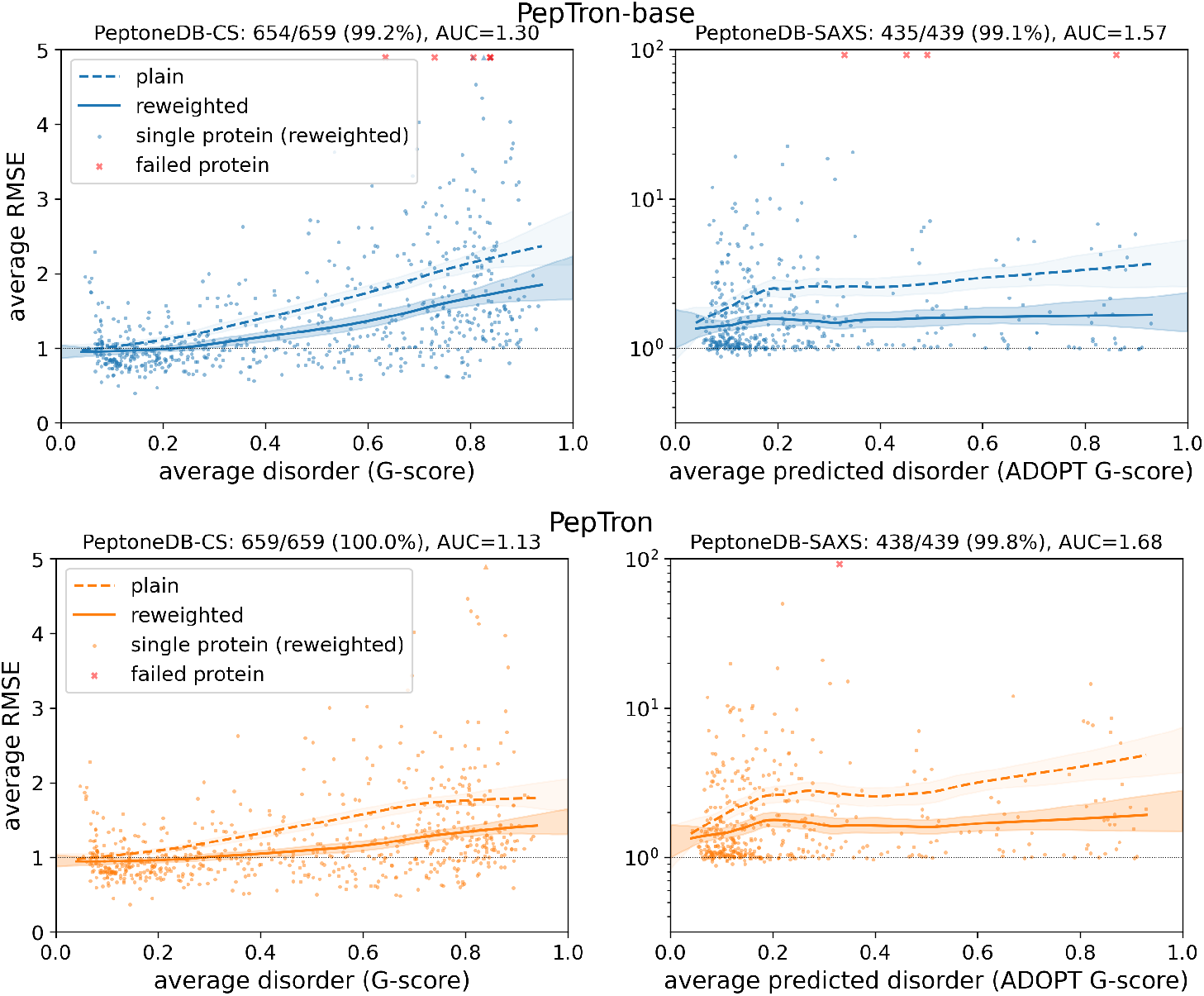
PeptoneBench results for the PepTron model before (PepTron-base) and after (PepTron) the fine-tuning on IDRome-o.

**Fig. D13.**
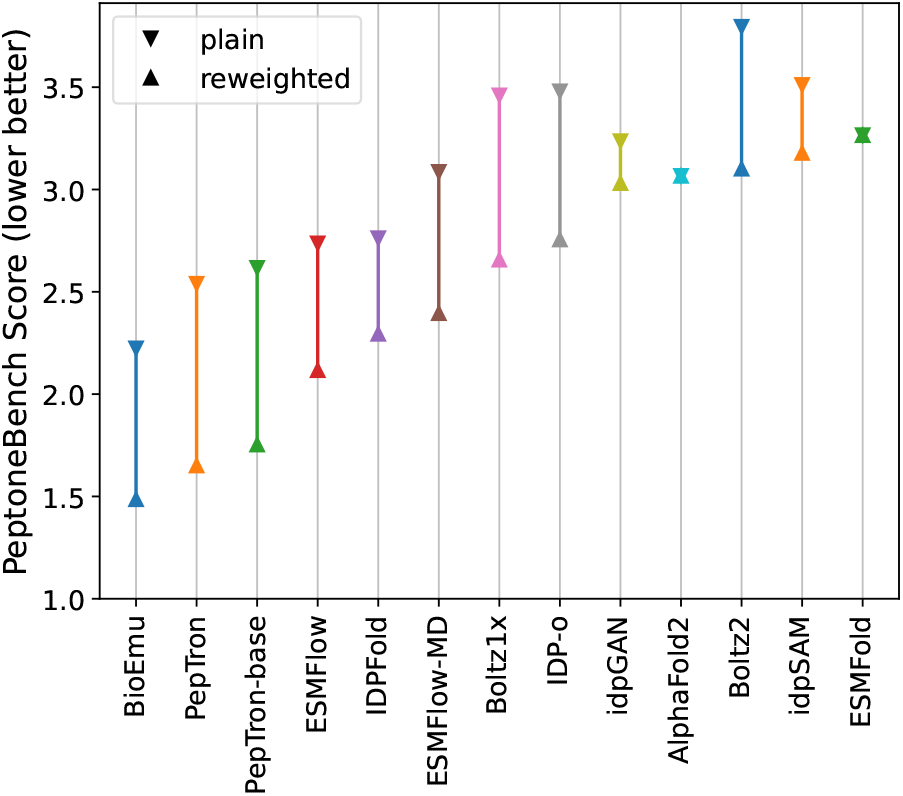
PeptoneBench summary score (the lower the better) for all the models tested, numerical values are in Table D3. The score is calculated according to Eq. D12. For each model, a line connecting the reweighted score (lower) and the plain score is shown. The longer the line, the more “reweightable” the model is.

**Table D3.** PeptoneBench summary score (the lower the better) for all the models tested, same data as Fig. D13. The score is calculated according to Eq. D12. The failed fraction *f* is obtained by normalizing the number of failed generations over the size of the respective dataset.

| Model | Score Reweighted | Score Plain | Failed Faction |
| --- | --- | --- | --- |
| BioEmu | 1.49 | 2.22 | 1.21% |
| PepTron | 1.65 | 2.54 | 0.11% |
| PepTron-base | 1.75 | 2.62 | 0.83% |
| ESMFlow | 2.12 | 2.74 | 1.67% |
| IDPFold | 2.29 | 2.76 | 23.99% |
| ESMFlow-MD | 2.40 | 3.09 | 0.11% |
| Boltz1x | 2.66 | 3.46 | 1.48% |
| IDP-o | 2.76 | 3.48 | 0.30% |
| IdpGAN | 3.03 | 3.24 | 35.15% |
| AlphaFold2 | 3.07 | 3.07 | 6.03% |
| Boltz2 | 3.10 | 3.80 | 0.27% |
| IdpSAM | 3.18 | 3.51 | 72.91% |
| ESMFold | 3.27 | 3.27 | 7.55% |

**Fig. D14.**
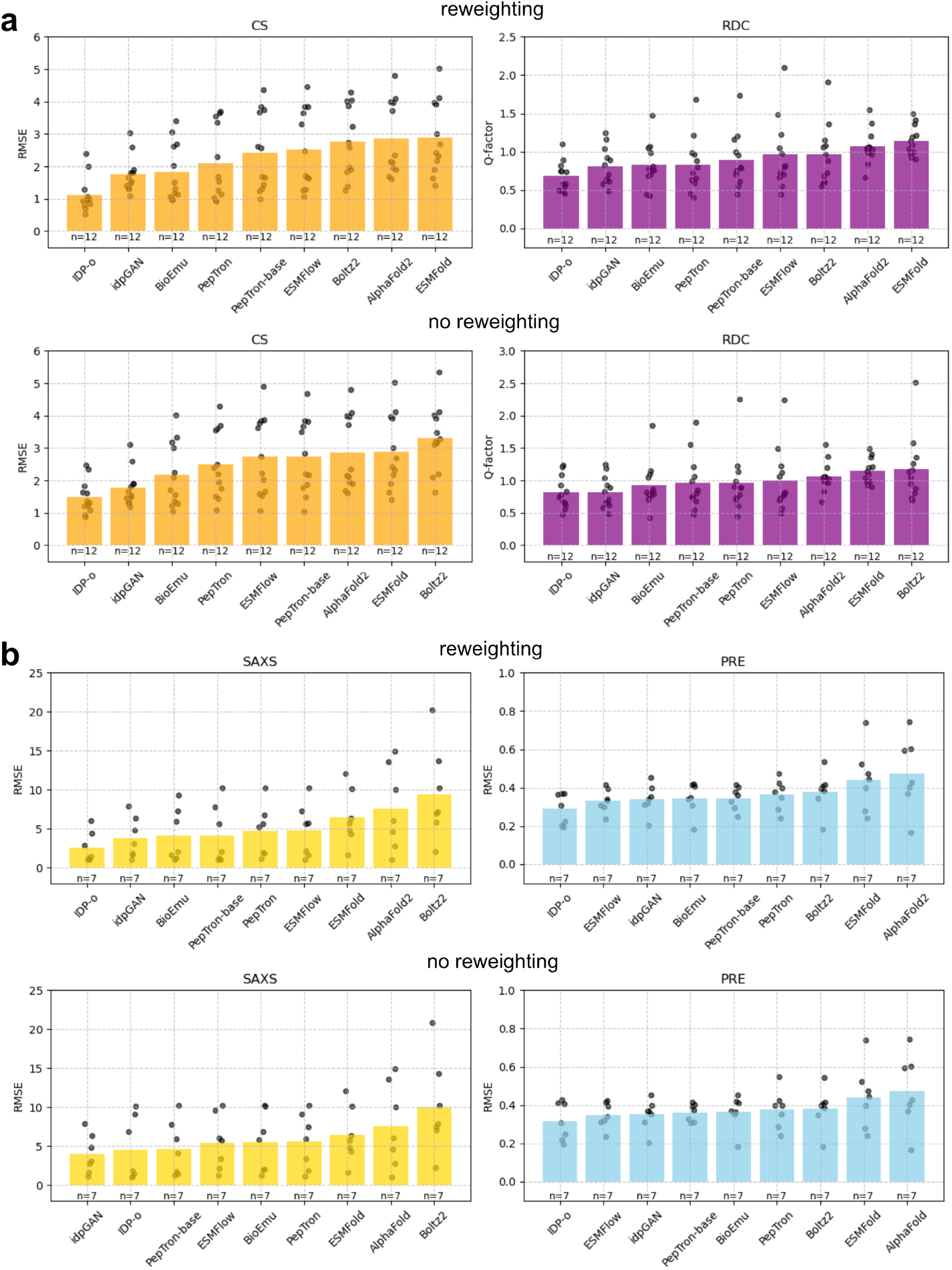
(a) RMSE between predicted and experimental chemical shifts and RDC Q-factors before and after maximum entropy reweighting with CS data. Each bar represents mean performance for n=12 intrinsically disordered systems for which both CS and RDC are available; individual points correspond to results for each individual system. Lower RMSE and Q-factor values indicate improved agreement with experimental data. Models are ranked from left to right for each metric. (b) RMSE between predicted and experimental SAXS profiles and PRE data before and after maximum entropy reweighting. Each bar represents mean performance for n=7 intrinsically disordered systems for which both SAXS and PREs are available. Models are ranked from left to right for each metric.

**Fig. D15.**
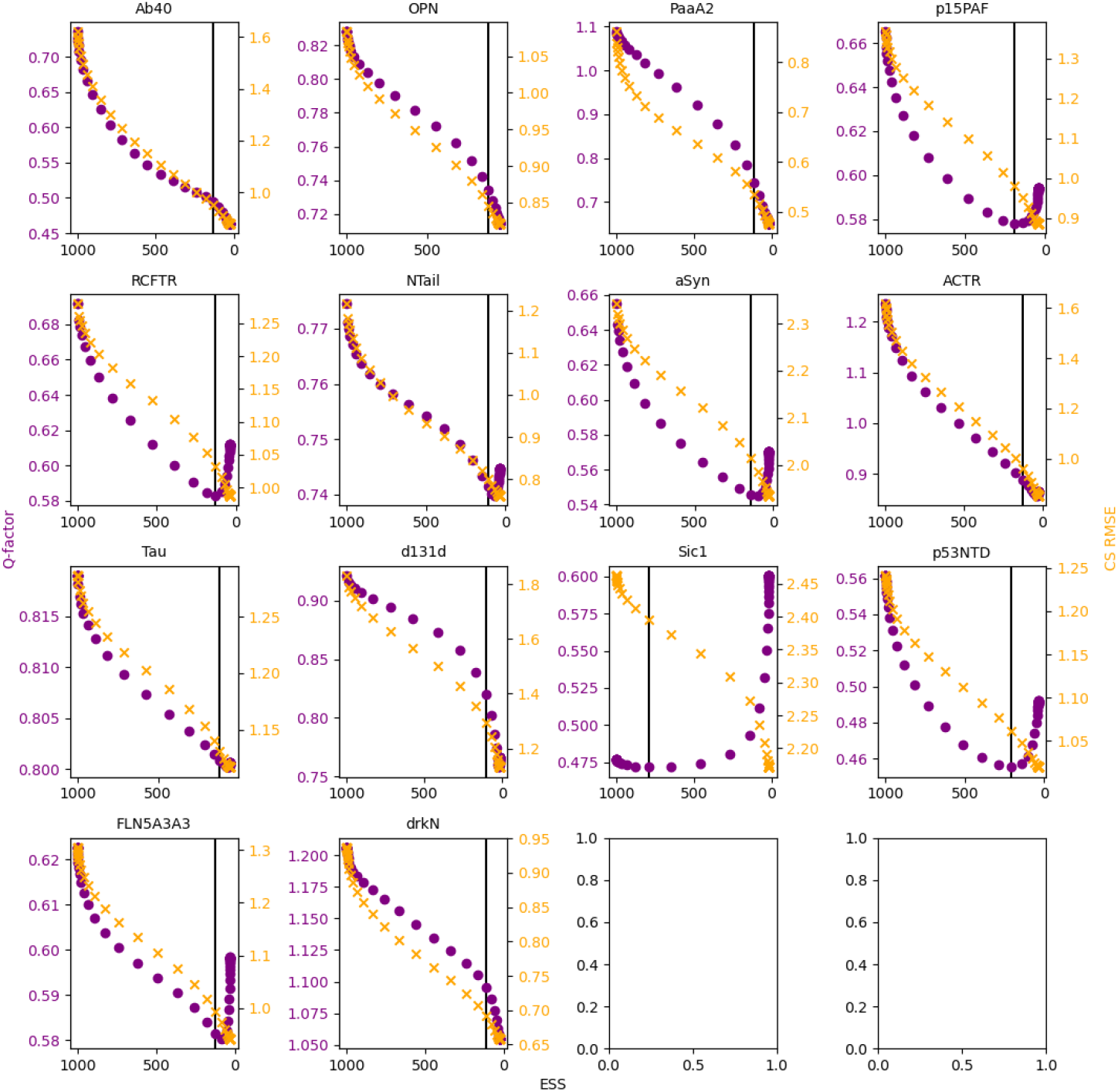
Example cross-correlation analyses of RDC Q-factors during maximum entropy reweighting with chemical shifts for all proteins predicted with the IDP-o model. The selected posterior ensemble is indicated by a black, vertical line. The x-axes depict the effective sample sizes (ESS).

**Fig. D16.**
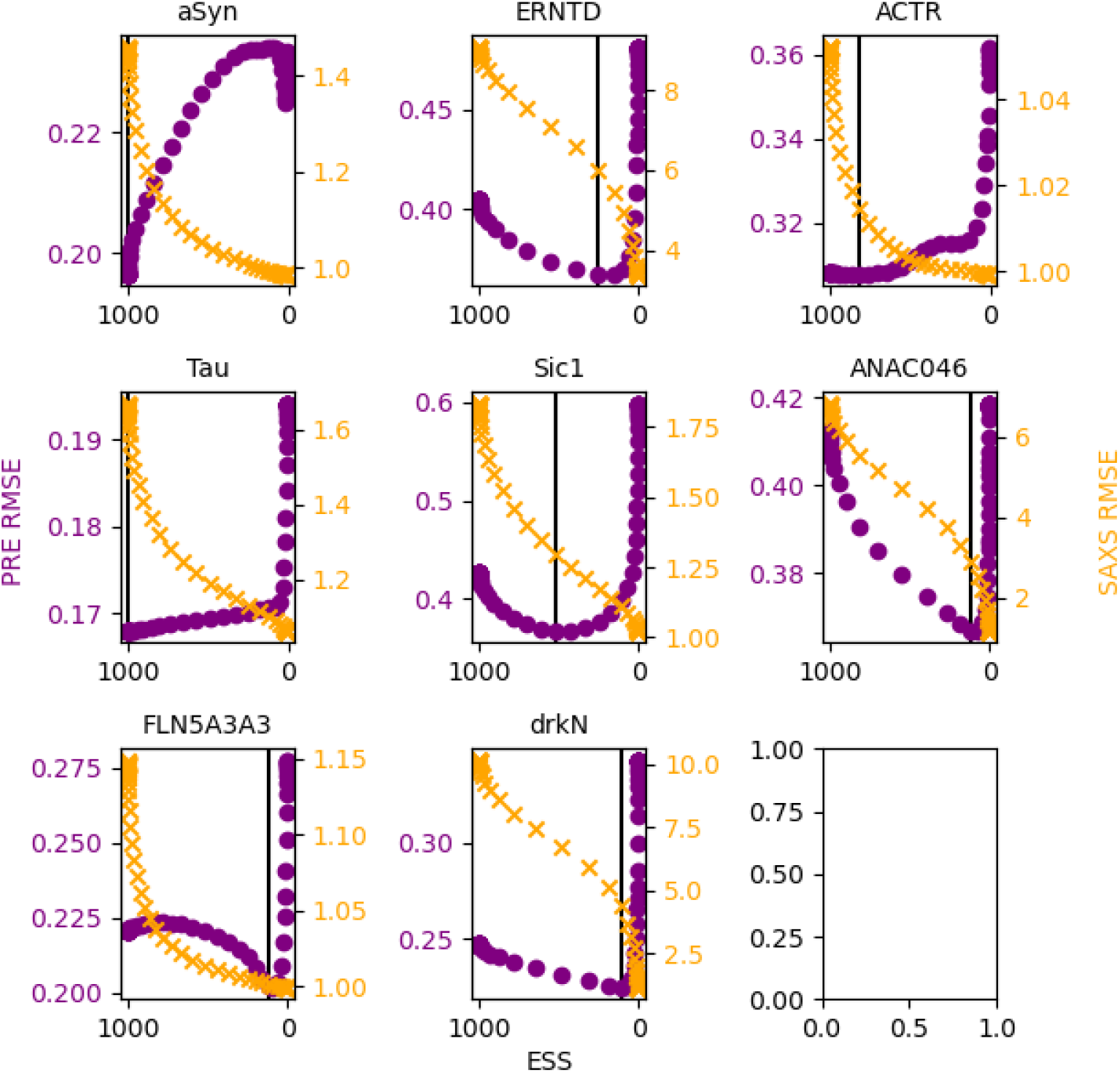
Example cross-correlation analyses of PRE RMSEs during maximum entropy reweighting with SAXS data for all proteins predicted with the IDP-o model. The selected posterior ensemble is indicated by a black, vertical line. The x-axes depict the effective sample sizes (ESS).

**Fig. D17.**
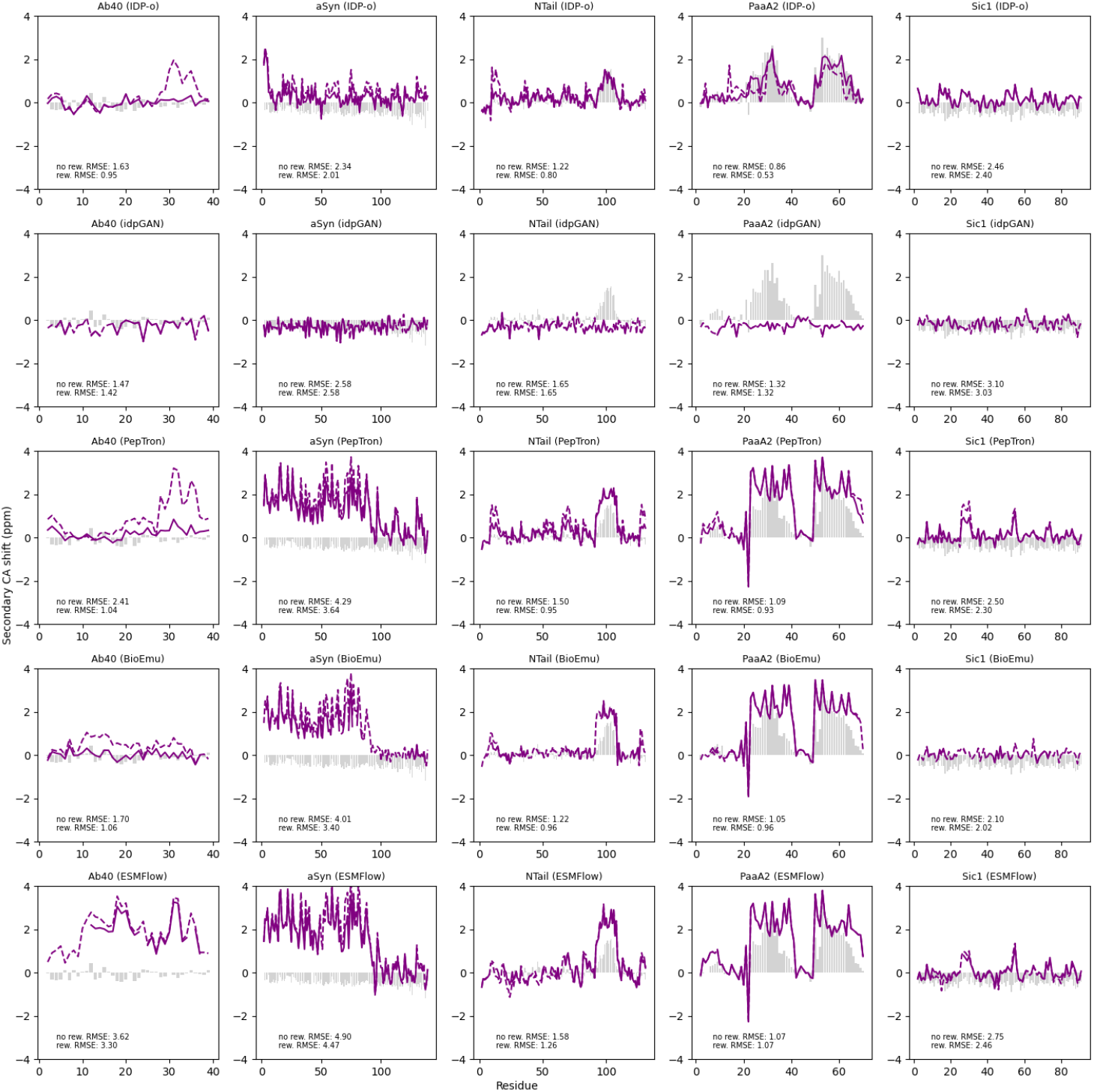
Predicted (purple lines) and experimental secondary CA chemical shifts (grey bars) for a set of exemplar proteins and models. Predicted values are depicted before (dashed lines) and after maximum entropy reweighting (solid lines). Reported RMSE values for each protein and model refer to the agreement with chemical shifts from all available nuclei.

**Fig. D18.**
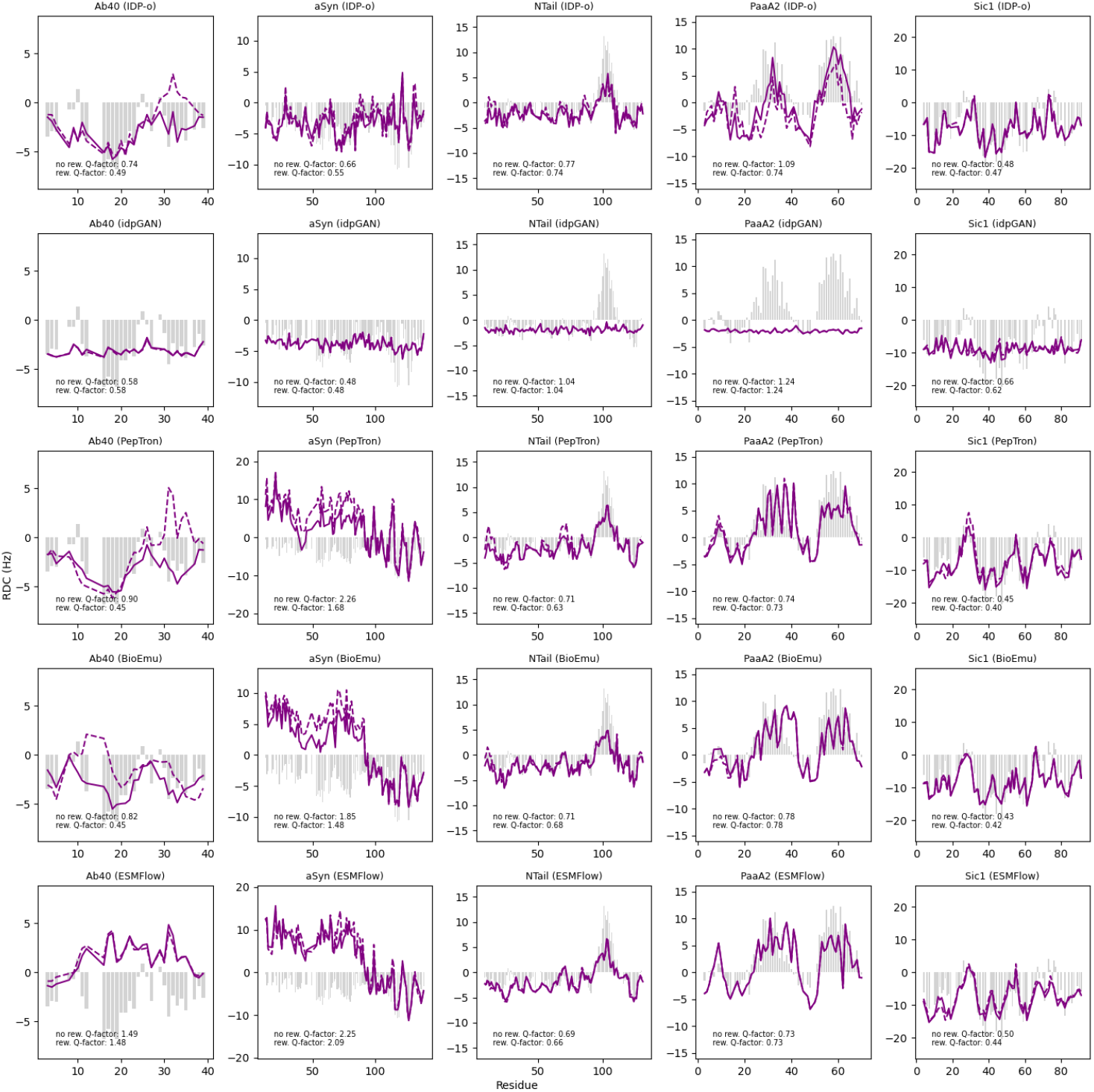
Predicted (purple lines) and experimental RDCs (grey bars) for a set of exemplar proteins and models. Predicted values are depicted before (dashed lines) and after maximum entropy reweighting (solid lines).

**Fig. D19.**
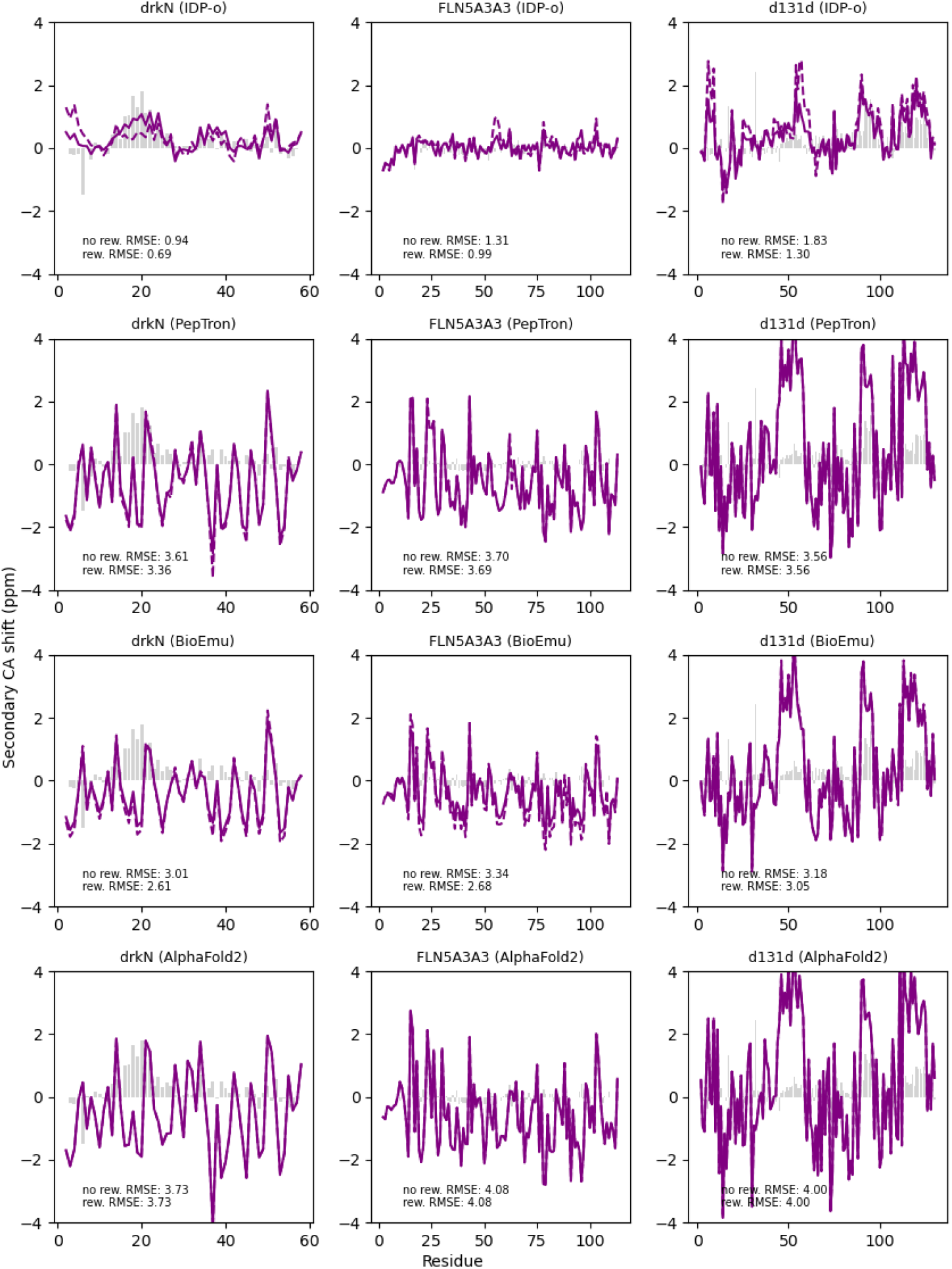
Predicted (purple lines) and experimental secondary CA chemical shifts (grey bars) for unfolded states of destabilized proteins with IDP-o, PepTron, BioEmu, and AlphaFold2. Predicted values are depicted before (dashed lines) and after maximum entropy reweighting (solid lines). Reported RMSE values for each protein and model refer to the agreement with chemical shifts from all available nuclei.

## Appendix E Other disorder metrics

Similar to the G-score, the Z-score [36] is derived from chemical shifts and spans a range from approximately -2 (fully disordered) to 16 (fully ordered). As previously shown, ADOPT [121] can predict accurately predict Z-scores from sequences. Figure E20 and E21 show the same benchmark figures as 2 and 3 but using predicted Z-scores instead of predicted G-scores.

**Fig. E20.**
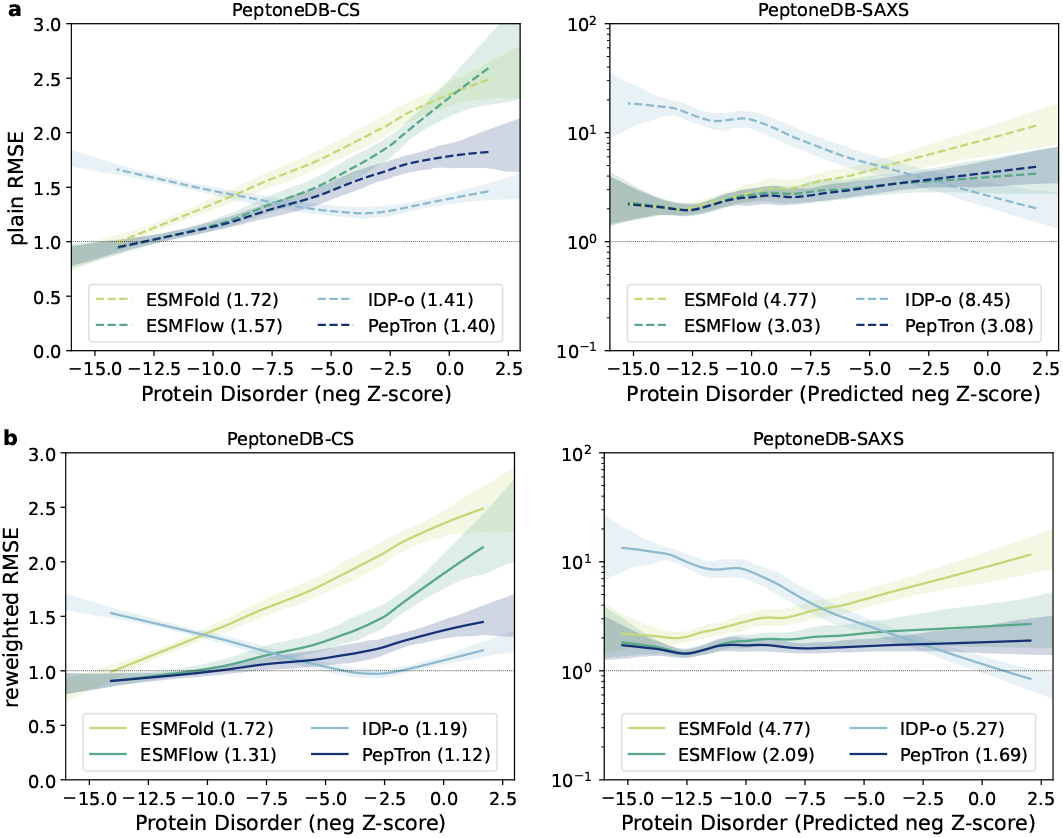
Same as Fig. 2 but using the negative Z-score instead of the G-score, and ADOPT instead of ADOPT2

**Fig. E21.**
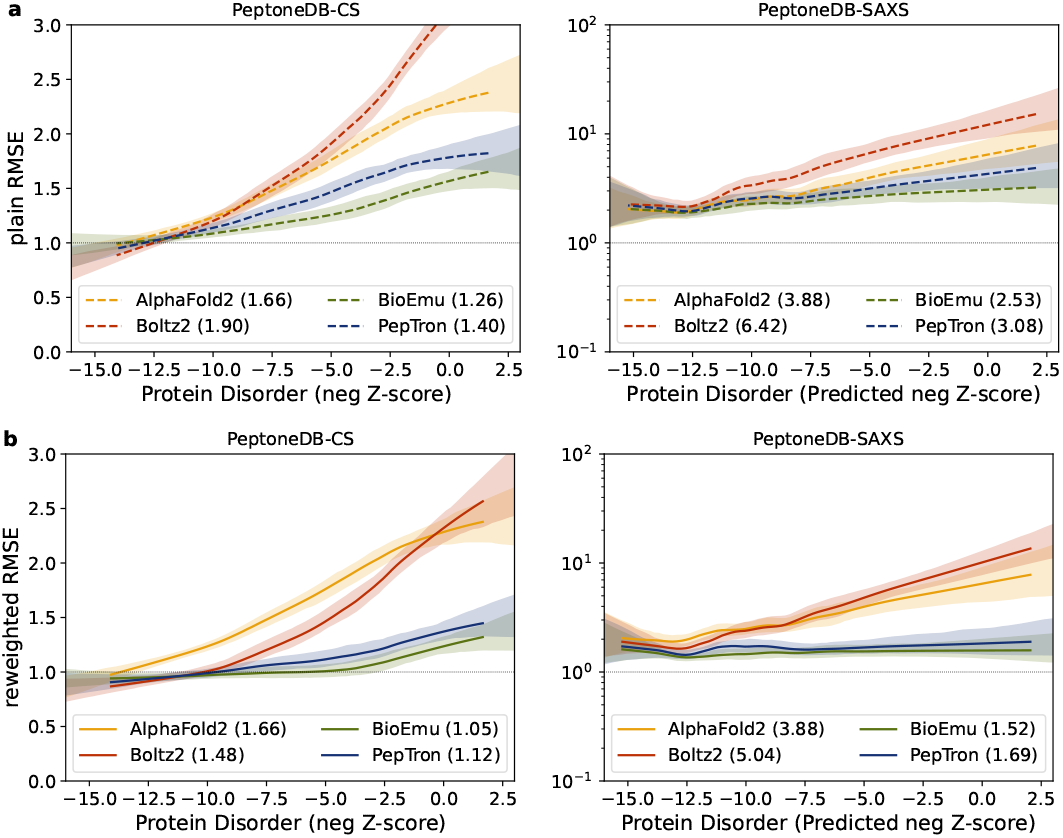
Same as Fig. 3 but using the negative Z-score instead of the G-score, and ADOPT instead of ADOPT2

## Footnotes

1 esm2nv-3B model from https://docs.nvidia.com/bionemo-framework/2.1/models/esm2

2 https://github.com/NVIDIA/bionemo-framework

3 https://docs.nvidia.com/cuda/cuequivariance/api/generated/cuequivariance_torch.triangle_attention.html

4 https://docs.nvidia.com/bionemo-framework/2.7rc1/main/developer-guide/bionemo-moco/bionemo-moco-Overview/

5 https://www.statsmodels.org/stable/generated/statsmodels.nonparametric.smoothers_lowess.lowess.html

6 https://github.com/biochunan/esmfold-docker-image/tree/5aaa4c0749c1875c50a8abd099849e78ee72d76c

7 https://github.com/feiglab/idpgan/tree/a414f3d50f6706ff4472ff83ff54505c8468bbaa

8 https://github.com/huhlim/cg2all/tree/v1.2

9 https://github.com/giacomo-janson/idpsam

10 See footnote 8

11 https://github.com/Junjie-Zhu/IDPFold/tree/ba40f40c3542b307c75d51b32665de23f89b4c47

12 https://github.com/microsoft/bioemu/tree/v.1.1.0

13 https://github.com/bjing2016/alphaflow

14 https://storage.googleapis.com/alphaflow/params/esmflow_pdb_base_202402.pt

15 See footnote 13

16 https://storage.googleapis.com/alphaflow/params/esmflow_md_base_202402.pt

17 https://github.com/microsoft/bioemu/tree/v.1.1.0

18 https://github.com/soedinglab/MMseqs2-App/tree/14e087560f309f989a5e1feb54fd1f9c988076d5

19 https://github.com/jwohlwend/boltz/tree/v1.0.0

20 See footnote 18

21 https://github.com/jwohlwend/boltz/tree/v2.1.0

22 See footnote 18

## References

[1] Holehouse, A. S. & Kragelund, B. B. The molecular basis for cellular function of intrinsically disordered protein regions. Nature Reviews Molecular Cell Biology 25, 187–211 (2024).

[2] Pritišanac, I. et al. A functional map of the human intrinsically disordered proteome. biorxiv [preprint](2024) (2025).

[3] Berman, H. M. et al. The protein data bank. Nucleic acids research 28, 235–242 (2000).

[4] Burley, S. K. et al. Updated resources for exploring experimentally-determined pdb structures and computed structure models at the rcsb protein data bank. Nucleic acids research 53, D564–D574 (2025).

[5] Gall, T. L., Romero, P. R., Cortese, M. S., Uversky, V. N. & Dunker, A. K. Intrinsic disorder in the protein data bank. Journal of Biomolecular Structure and Dynamics 24, 325–341 (2007).

[6] Erdős, G. & Dosztanyi, Z. Deep learning for intrinsically disordered proteins: From improved predictions to deciphering conformational ensembles. Current Opinion in Structural Biology 89, 102950 (2024).

[7] Lotthammer, J. M., Ginell, G. M., Griffith, D., Emenecker, R. & Holehouse, A. S. Direct prediction of intrinsically disordered protein conformational properties from sequence. Biophysical Journal 123, 43a (2024).

[8] Riley, A. C., Ashlock, D. A. & Graether, S. P. The difficulty of aligning intrinsically disordered protein sequences as assessed by conservation and phylogeny. Plos one 18, e0288388 (2023).

[9] Jumper, J. et al. Highly accurate protein structure prediction with alphafold. nature 596, 583–589 (2021).

[10] Piovesan, D., Monzon, A. M. & Tosatto, S. C. Intrinsic protein disorder and conditional folding in alphafolddb. Protein Science 31, e4466 (2022).

[11] Abramson, J. et al. Accurate structure prediction of biomolecular interactions with alphafold 3. Nature 630, 493–500 (2024).

[12] Corley, N. et al. Accelerating biomolecular modeling with atomworks and rf3. bioRxiv 2025–08 (2025).

[13] Jing, B., Berger, B. & Jaakkola, T. AlphaFold meets flow matching for generating protein ensembles. NeurIPS (2023). URL https://openreview.net/forum?id=yQcebEgQfH. Three versions, only the first one mentions IDPs.

[14] Lewis, S. et al. Scalable emulation of protein equilibrium ensembles with generative deep learning. Science eadv9817 (2025).

[15] Cao, F., von Bülow, S., Tesei, G. & Lindorff-Larsen, K. A coarse-grained model for disordered and multidomain proteins. Protein Science 33, e5172 (2024).

[16] Joseph, J. A. et al. Physics-driven coarse-grained model for biomolecular phase separation with near-quantitative accuracy. Nature computational science 1, 732–743 (2021).

[17] Janson, G. & Feig, M. Transferable deep generative modeling of intrinsically disordered protein conformations. PLOS Computational Biology 20, e1012144 (2024).

[18] Janson, G., Valdes-Garcia, G., Heo, L. & Feig, M. Direct generation of protein conformational ensembles via machine learning. Nature Communications 14, 774 (2023).

[19] Zhang, O., Liu, Z. H., Forman-Kay, J. D. & Head-Gordon, T. Deep learning of proteins with local and global regions of disorder. arXiv preprint 2502.11326 (2025).

[20] Schnapka, V., Morozova, T. I., Sen, S. & Bonomi, M. Atomic resolution ensembles of intrinsically disordered proteins with alphafold. bioRxiv 2025.06.18.660298 (2025).

[21] von Bülow, S., Johansson, K. E. & Lindorff-Larsen, K. Af-calvados: Alphafold-guided simulations of multi-domain proteins at the proteome level. bioRxiv 2025–10 (2025).

[22] Ghafouri, H. et al. Ped in 2024: improving the community deposition of structural ensembles for intrinsically disordered proteins. Nucleic acids research 52, D536–D544 (2024).

[23] Sil, S., Datta, I. & Basu, S. Use of ai-methods over md simulations in the sampling of conformational ensembles in idps. Frontiers in Molecular Biosciences 12, 1542267 (2025).

[24] Ghafouri, H. et al. Towards a unified framework for determining conformational ensembles of disordered proteins. arXiv (2025).

[25] Norgaard, A. B., Ferkinghoff-Borg, J. & Lindorff-Larsen, K. Experimental parameterization of an energy function for the simulation of unfolded proteins. Biophysical journal 94, 182–192 (2008).

[26] Robustelli, P., Piana, S. & Shaw, D. E. Developing a molecular dynamics force field for both folded and disordered protein states. Proceedings of the National Academy of Sciences 115, E4758–E4766 (2018).

[27] Borthakur, K., Sisk, T. R., Panei, F. P., Bonomi, M. & Robustelli, P. Determining accurate conformational ensembles of intrinsically disordered proteins at atomic resolution. Nature Communications 16, 9036 (2025). URL 10.1038/s41467-025-64098-3.

[28] Hummer, G. & Köfinger, J. Bayesian ensemble refinement by replica simulations and reweighting. The Journal of chemical physics 143 (2015).

[29] Cesari, A., Gil-Ley, A. & Bussi, G. Combining simulations and solution experiments as a paradigm for rna force field refinement. Journal of chemical theory and computation 12, 6192–6200 (2016).

[30] Bonomi, M., Camilloni, C., Cavalli, A. & Vendruscolo, M. Metainference: A bayesian inference method for heterogeneous systems. Science advances 2, e1501177 (2016).

[31] Bottaro, S., Bengtsen, T. & Lindorff-Larsen, K. Integrating Molecular Simulation and Experimental Data: A Bayesian/Maximum Entropy Reweighting Approach, 219–240 (Springer US, New York, NY, 2020).

[32] Xue, B., Dunker, A. K. & Uversky, V. N. Orderly order in protein intrinsic disorder distribution: disorder in 3500 proteomes from viruses and the three domains of life. Journal of Biomolecular Structure and Dynamics 30, 137–149 (2012).

[33] Aspromonte, M. C. et al. Disprot in 2024: improving function annotation of intrinsically disordered proteins. Nucleic Acids Research 52, D434–D441 (2024).

[34] Tesei, G. et al. Conformational ensembles of the human intrinsically disordered proteome. Nature 626, 897–904 (2024).

[35] Haak, M., Senoner, T., Schlensok, J., Wagemann, D. & Rost, B. TriZOD: continuous, per-residue disorder scores from protein NMR experiments. https://github.com/MarkusHaak/trizod (2024).

[36] Nielsen, J. T. & Mulder, F. A. A. Quality and bias of protein disorder predictors. Scientific Reports 9, 5137 (2019).

[37] Redl, I. et al. Optimizing protein language models with sentence transformers (2023). URL https://www.mlsb.io/papers 2023/Optimizing protein language models with Sentence Transformers.pdf. Work-shop Paper.

[38] Hoch, J. C. et al. Biological magnetic resonance data bank. Nucleic Acids Research 51, D368–D376 (2022).

[39] Kikhney, A. G., Borges, C. R., Molodenskiy, D. S., Jeffries, C. M. & Svergun, D. I. SASBDB: Towards an automatically curated and validated repository for biological scattering data. Protein Science 29, 66–75 (2020).

[40] Thomasen, F. E. et al. Rescaling protein-protein interactions improves martini 3 for flexible proteins in solution. Nature Communications 15, 6645 (2024).

[41] Cavender, C. E. et al. Structure-based experimental datasets for benchmarking of protein simulation force fields. arXiv preprint 2303.11056 (2023).

[42] Shen, Y. & Bax, A. SPARTA+: a modest improvement in empirical NMR chemical shift prediction by means of an artificial neural network. Journal of Biomolecular NMR 48, 13–22 (2010).

[43] Li, J., Bennett, K. C., Liu, Y., Martin, M. V. & Head-Gordon, T. Accurate prediction of chemical shifts for aqueous protein structure on “real world” data. Chemical Science 11, 3180–3191 (2020).

[44] Grudinin, S., Garkavenko, M. & Kazennov, A. Pepsi-SAXS: an adaptive method for rapid and accurate computation of small-angle x-ray scattering profiles. Acta Crystallographica Section D: Structural Biology 73, 449–464 (2017).

[45] Sterckx, Y. G. et al. Small-angle x-ray scattering-and nuclear magnetic resonance-derived conformational ensemble of the highly flexible antitoxin paaa2. Structure 22, 854–865 (2014).

[46] Tesei, G. et al. Deer-predict: Software for efficient calculation of spin-labeling epr and nmr data from conformational ensembles. PLOS Computational Biology 17, e1008551 (2021).

[47] Teixeira, J. M. et al. Idpconformergenerator: a flexible software suite for sampling the conformational space of disordered protein states. The Journal of Physical Chemistry A 126, 5985–6003 (2022).

[48] Pietrek, L. M., Stelzl, L. S. & Hummer, G. Hierarchical ensembles of intrinsically disordered proteins at atomic resolution in molecular dynamics simulations. Journal of chemical theory and computation 16, 725–737 (2019).

[49] John, P. S. et al. BioNeMo framework: a modular, high-performance library for AI model development in drug discovery. arXiv (2024).

[50] Shoeybi, M. et al. Megatron-lm: Training multi-billion parameter language models using model parallelism. arXiv preprint 1909.08053 (2019).

[51] Lin, Z. et al. Language models of protein sequences at the scale of evolution enable accurate structure prediction. bioRxiv (2022).

[52] Lin, Z. et al. Evolutionary-scale prediction of atomic-level protein structure with a language model. Science 379, 1123–1130 (2023).

[53] Passaro, S. et al. Boltz-2: Towards accurate and efficient binding affinity prediction. BioRxiv 2025–06 (2025).

[54] Wang, Y., Lu, J., Jaitly, N., Susskind, J. & Bautista, M. A. Simplefold: Folding proteins is simpler than you think. arXiv preprint 2509.18480 (2025).

[55] Invernizzi, M., Piaggi, P. M. & Parrinello, M. Unified approach to enhanced sampling. Physical Review X 10, 041034 (2020).

[56] Vander Meersche, Y., Cretin, G., Gheeraert, A.Gelly, J.-C. & Galochkina, T. Atlas: protein flexibility description from atomistic molecular dynamics simulations. Nucleic acids research 52, D384–D392 (2024).

[57] Mirarchi, A., Giorgino, T. & De Fabritiis, G. mdcath: A large-scale md dataset for data-driven computational biophysics. Scientific Data 11, 1299 (2024).

[58] Mokhtari, O., Bignon, E., Khakzad, H. & Karami, Y. Dynarepo: The repository of macromolecular conformational dynamics. bioRxiv 2025–08 (2025).

[59] Siebenmorgen, T. et al. Misato: machine learning dataset of protein–ligand complexes for structure-based drug discovery. Nature computational science 4, 367–378 (2024).

[60] Lindorff-Larsen, K. & Kragelund, B. B. On the potential of machine learning to examine the relationship between sequence, structure, dynamics and function of intrinsically disordered proteins. Journal of Molecular Biology 433, 167196 (2021).

[61] Schneidman-Duhovny, D., Pellarin, R. & Sali, A. Uncertainty in integrative structural modeling. Current opinion in structural biology 28, 96–104 (2014).

[62] Nagy, G., Hoffmann, S. V., Jones, N. C. & Grubmüller, H. Reference data set for circular dichroism spectroscopy comprised of validated intrinsically disordered protein models. Applied Spectroscopy 78, 897– 911 (2024).

[63] Wu, K. et al. Design of intrinsically disordered region binding proteins. Science 389, eadr8063 (2025).

[64] Amaro, R. E. et al. The need to implement fair principles in biomolecular simulations. Nature methods 1–5 (2025).

[65] Heo, L. & Feig, M. One particle per residue is sufficient to describe all-atom protein structures. bioRxiv (2023).

[66] Visani, G. M., Galvin, W., Pun, M. & Nourmohammad, A. Knowles, D. A. & Mostafavi, S. (eds) H-packer: Holographic rotationally equivariant convolutional neural network for protein side-chain packing. (eds Knowles, D.A. & Mostafavi, S.) Proceedings of the 18th Machine Learning in Computational Biology meeting, Vol. 240 of Proceedings of Machine Learning Research, 230–249 (PMLR, 2024).

[67] Ptaszek, A. L., Li, J., Konrat, R., Platzer, G. & Head-Gordon, T. UCBShift 2.0: Bridging the gap from backbone to side chain protein chemical shift prediction for protein structures. Journal of the American Chemical Society 146, 31733–31745 (2024).

[68] Cavender, C. E. et al. Structure-based experimental datasets for benchmarking of protein simulation force fields. arXiv (2023).

[69] Nielsen, J. T. & Mulder, F. A. A. POTENCI: prediction of temperature, neighbor and pH-corrected chemical shifts for intrinsically disordered proteins. Journal of Biomolecular NMR 70, 141–165 (2018).

[70] Svergun, D. I. Determination of the regularization parameter in indirect-transform methods using perceptual criteria. Journal of Applied Crystallography 25, 495–503 (1992).

[71] Hansen, S. Bayesian estimation of hyperparameters for indirect fourier transformation in small-angle scattering. Applied Crystallography 33, 1415–1421 (2000).

[72] Steinegger, M. & Söding, J. MMseqs2 enables sensitive protein sequence searching for the analysis of massive data sets. Nature Biotechnology 35, 1026–1028 (2017).

[73] Pesce, F. & Lindorff-Larsen, K. Refining conformational ensembles of flexible proteins against small-angle x-ray scattering data. Biophysical Journal 120, 5124–5135 (2021).

[74] Hou, L. et al. Solution nmr studies of the aβ (1-40) and aβ (1-42) peptides establish that the met35 oxidation state affects the mechanism of amyloid formation. Journal of the American Chemical Society 126, 1992–2005 (2004).

[75] Newby, F. N. et al. Structure-free validation of residual dipolar coupling and paramagnetic relaxation enhancement measurements of disordered proteins. Biochemistry 54, 6876–6886 (2015).

[76] Iešmantavičius, V. et al. Modulation of the intrinsic helix propensity of an intrinsically disordered protein reveals long-range helix–helix interactions. Journal of the American Chemical Society 135, 10155–10163 (2013).

[77] Kjaergaard, M. et al. Temperature-dependent structural changes in intrinsically disordered proteins: Formation of α–helices or loss of polyproline ii? Protein Science 19, 1555–1564 (2010).

[78] Newcombe, E. A. et al. Insight into calcium-binding motifs of intrinsically disordered proteins. Biomolecules 11, 1173 (2021).

[79] Pesce, F. et al. Assessment of models for calculating the hydrodynamic radius of intrinsically disordered proteins. Biophysical Journal 122, 310–321 (2023).

[80] Due, A. D. et al. Hierarchy in regulator interactions with distant transcriptional activation domains empowers rheostatic regulation. Protein Science 34, e70142 (2025).

[81] Rao, J. N., Kim, Y. E., Park, L. S. & Ulmer, T. S. Effect of pseudorepeat rearrangement on α-synuclein misfolding, vesicle binding, and micelle binding. Journal of molecular biology 390, 516–529 (2009).

[82] Bertoncini, C. W. et al. Release of long-range tertiary interactions potentiates aggregation of natively unstructured α-synuclein. Proceedings of the National Academy of Sciences 102, 1430–1435 (2005).

[83] Ahmed, M. C. et al. Refinement of α-synuclein ensembles against saxs data: Comparison of force fields and methods. Frontiers in molecular biosciences 8, 654333 (2021).

[84] Dedmon, M. M., Lindorff-Larsen, K., Christodoulou, J., Vendruscolo, M. & Dobson, C. M. Mapping long-range interactions in α-synuclein using spin-label nmr and ensemble molecular dynamics simulations. Journal of the American Chemical Society 127, 476–477 (2005).

[85] Qu, X. et al. Structural basis for the dynamic chaperoning of disordered clients by hsp90. Nature Structural & Molecular Biology 31, 1482–1491 (2024).

[86] Shortle, D. & Ackerman, M. S. Persistence of native-like topology in a denatured protein in 8 m urea. Science 293, 487–489 (2001).

[87] Zhang, O. & Forman-Kay, J. D. Structural characterization of folded and unfolded states of an sh3 domain in equilibrium in aqueous buffer. Biochemistry 34, 6784–6794 (1995).

[88] Marsh, J. A., Baker, J. M., Tollinger, M. & Forman-Kay, J. D. Calculation of residual dipolar couplings from disordered state ensembles using local alignment. Journal of the American Chemical Society 130, 7804–7805 (2008).

[89] Choy, W.-Y. et al. Distribution of molecular size within an unfolded state ensemble using small-angle x-ray scattering and pulse field gradient nmr techniques. Journal of molecular biology 316, 101–112 (2002).

[90] Marsh, J. A. & Forman-Kay, J. D. Structure and disorder in an unfolded state under nondenaturing conditions from ensemble models consistent with a large number of experimental restraints. Journal of molecular biology 391, 359–374 (2009).

[91] Du, Z. et al. The sequence–structure–function relationship of intrinsic erα disorder. Nature 638, 1130–1138 (2025).

[92] Streit, J. O. et al. The ribosome lowers the entropic penalty of protein folding. Nature 633, 232–239 (2024).

[93] Gely, S. et al. Solution structure of the c-terminal x domain of the measles virus phosphoprotein and interaction with the intrinsically disordered c-terminal domain of the nucleoprotein. Journal of molecular recognition 23, 435–447 (2010).

[94] Jensen, M. R. et al. Intrinsic disorder in measles virus nucleocapsids. Proceedings of the National Academy of Sciences 108, 9839–9844 (2011).

[95] Schedlbauer, A. et al. Backbone assignment of osteopontin, a cytokine and cell attachment protein implicated in tumorigenesis. Biomolecular NMR Assignments 2, 29–31 (2008).

[96] Platzer, G. et al. The metastasis-associated extracellular matrix protein osteopontin forms transient structure in ligand interaction sites. Biochemistry 50, 6113–6124 (2011).

[97] De Biasio, A. et al. p15paf is an intrinsically disordered protein with nonrandom structural preferences at sites of interaction with other proteins. Biophysical journal 106, 865–874 (2014).

[98] Sebák, F., Ecsédi, P., Nyitray, L. & Bodor, A. Assignment of the disordered, proline-rich n-terminal domain of the tumour suppressor p53 protein using 1hn and 1hα-detected nmr measurements. Biomolecular NMR Assignments 17, 309–314 (2023).

[99] Wells, M. et al. Structure of tumor suppressor p53 and its intrinsically disordered n-terminal transactivation domain. Proceedings of the National academy of Sciences 105, 5762–5767 (2008).

[100] Baker, J. M. et al. Cftr regulatory region interacts with nbd1 predominantly via multiple transient helices. Nature structural & molecular biology 14, 738–745 (2007).

[101] Mittag, T. et al. Structure/function implications in a dynamic complex of the intrinsically disordered sic1 with the cdc4 subunit of an scf ubiquitin ligase. Structure 18, 494–506 (2010).

[102] Gomes, G.-N. W. et al. Conformational ensembles of an intrinsically disordered protein consistent with nmr, saxs, and single-molecule fret. Journal of the American Chemical Society 142, 15697–15710 (2020).

[103] Crha, R. et al. Hiding in plain sight: complex interaction patterns between tau and 14-3-3? protein variants. International Journal of Biological Macromolecules 266, 130802 (2024).

[104] Mukrasch, M. D. et al. Structural polymorphism of 441-residue tau at single residue resolution. PLoS biology 76, e1000034 (2009).

[105] Mylonas, E. et al. Domain conformation of tau protein studied by solution small-angle x-ray scattering. Biochemistry 47, 10345–10353 (2008).

[106] Zweckstetter, M. Nmr: prediction of molecular alignment from structure using the pales software. Nature protocols 3, 679–690 (2008).

[107] Kim, H., Mirdita, M. & Steinegger, M. Foldcomp: a library and format for compressing and indexing large protein structure sets. Bioinformatics 39, btad153 (2023).

[108] Dutton, O., Hoffmann, F. & Tamiola, K. Nerfax: An efficient and scalable conversion from the internal representation to cartesian space. bioRxiv 2022–05 (2022).

[109] Verkuil, R. et al. Language models generalize beyond natural proteins. BioRxiv 2022–12 (2022).

[110] Consortium, U. Uniprot: the universal protein knowledgebase in 2021. Nucleic acids research 49, D480–D489 (2021).

[111] Ahdritz, G. et al. Openfold: Retraining alphafold2 yields new insights into its learning mechanisms and capacity for generalization. Nature methods 21, 1514–1524 (2024).

[112] Webby, M. N. et al. Structural Analysis of the Menangle Virus P Protein Reveals a Soft Boundary between Ordered and Disordered Regions. Viruses 13, 1737 (2021).

[113] Robin, X. et al. Continuous automated model evaluation (cameo)—perspectives on the future of fully automated evaluation of structure prediction methods. Proteins: Structure, Function, and Bioinformatics 89, 1977–1986 (2021).

[114] Jing, B. et al. Eigenfold: Generative protein structure prediction with diffusion models. arXiv preprint 2304.02198 (2023).

[115] Mirdita, M. et al. Colabfold: making protein folding accessible to all. Nature methods 19, 679–682 (2022).

[116] Lin, Z. et al. Evolutionary-scale prediction of atomic-level protein structure with a language model. Science 379, 1123–1130 (2023).

[117] Janson, G., Valdes-Garcia, G., Heo, L. & Feig, M. Direct generation of protein conformational ensembles via machine learning. Nature Communications 14, 774 (2023). idpGAN.

[118] Janson, G. & Feig, M. Transferable deep generative modeling of intrinsically disordered protein conformations. bioRxiv 2024.02.08.579522 (2024).

[119] Zhu, J. et al. Accurate Generation of Conformational Ensembles for Intrinsically Disordered Proteins with IDPFold. Advanced Science (2025).

[120] Wohlwend, J. et al. Boltz-1 democratizing biomolecular interaction modeling. bioRxiv 2024.11.19.624167 (2024).

[121] Redl, I. et al. ADOPT: intrinsic protein disorder prediction through deep bidirectional transformers. NAR Genomics and Bioinformatics 5 (2023). URL 10.1093/nargab/lqad041.Lqad041.

